# Identifying dysregulated immune cell subsets following critical volumetric muscle loss with pseudo-time trajectories

**DOI:** 10.1101/2021.05.25.445480

**Authors:** Lauren A. Hymel, Shannon E. Anderson, Thomas C. Turner, William Y. York, Hong Seo Lim, Peng Qiu, Young C. Jang, Nick J. Willett, Edward A. Botchwey

**Author notes:** Denotes co-first author (Lauren A. Hymel, Shannon E. Anderson).

## Abstract

Volumetric muscle loss (VML) results in permanent functional deficits and remains a substantial regenerative medicine challenge. A coordinated immune response is crucial for timely myofiber regeneration, however the immune response following VML has yet to be fully characterized. Here, we leveraged dimensionality reduction and pseudo-time analysis techniques to elucidate the cellular players underlying a functional or pathological outcome as a result of subcritical or critical VML in the murine quadriceps, respectively. We found that critical VML presented with a sustained presence of M2-like and CD206^hi^Ly6C^hi^ ‘hybrid’ macrophages whereas subcritical defects resolved these populations. These macrophage subsets may contribute to fibrogenesis in critical VML, especially in the presence of TGF-β. Furthermore, several T cell populations were significantly elevated in critical VML compared to subcritical injuries. Specifically, there was a significant increase of CD127^+^ T cells at days 3 and 7, and upregulated CD127 expression may indicate aberrant IL-7 signaling in critical VML. These results demonstrate a dysregulated immune response in critical VML that is unable to resolve the chronic inflammatory state and transition to a pro-regenerative microenvironment. These data provide important insights into potential therapeutic strategies which could reduce the immune cell burden and pro-fibrotic signaling characteristic of VML.

## Introduction

Following acute muscle injury, skeletal muscle’s robust regenerative response relies on the prompt and proper coordination of immune cells. The cellular dynamics of myeloid and lymphoid cell trafficking into and out of the muscle coincide with each stage of the muscle regenerative process (1). Following muscle injury, there is degeneration and necrosis of damaged myofibers (2), triggering the invasion of neutrophils that peak within hours following injury and drop off after 24 hours (3). As neutrophils secrete tumor necrosis factor (TNF) and interferon-γ (IFN-γ), monocyte derived pro-inflammatory phagocytic (M1) macrophages infiltrate the muscle to aid in the removal of tissue debris and propagate pro-inflammatory signals by secretion of cytokines (1). Both TNF-α and IFN-γ play a role in macrophage induction and skeletal muscle regeneration by silencing Pax7 and preventing the expression of MyoD to maintain a required pool of skeletal muscle stem/satellite cells (MuSCs), in an activated, proliferative state (1, 4). M1 macrophage secretion of TNF-α is also a mechanism to induce fibroadipogenic progenitor cell (FAP) clearance by apoptosis (5). FAPs are muscle-resident mesenchymal stromal cells, and their regulated apoptosis induced by TNF-α is a key signaling event for healthy extracellular matrix (ECM) secretion (6).

Once in tissue, classical Ly6C^hi^ monocytes can convert into non-classical Ly6C^lo^ monocytes which are biased progenitors of pro-regenerative (M2) macrophages, as we have shown previously (7). These pro-regenerative or alternatively activated M2 macrophages peak in concentration between 4-7 days following muscle injury (8, 9). The transition to an anti-inflammatory, interleukin (IL)-10 and transforming growth factor (TGF)-β rich environment corresponds with both a transition to M2 macrophages as well as the differentiation and growth stages of myogenesis (10). As the skeletal muscle regenerates, myeloid cells traffic out of the tissue by two weeks post-injury (11). This coordinated response of myeloid cells is crucial for the proper regeneration of skeletal muscle, as the depletion or altered polarization of macrophages has been shown to increase adipose and fibrotic tissue deposition while reducing regenerated myofiber cross-sectional area (12, 13).

In addition to myeloid derived cells, lymphoid-derived T-cells also respond to muscle injury. Peak infiltration of CD4^+^ helper and CD8^+^ cytotoxic T cells occurs at 3 days post-injury, returning to baseline levels gradually by day 14 (14). Regulatory T cells (T_reg_ cells) infiltrate the muscle after acute injury with similar kinetics to that of M2 macrophages – peaking at day 4 post-injury (15). The T cell response has been implicated in the maintenance of myeloid cell infiltration. The depletion of CD8^+^ T cells have been shown to reduce skeletal muscle regeneration through a reduction in the recruitment of pro-inflammatory M1 macrophages (14). Similarly, depletion of T_reg_ cells impairs muscle repair and prolongs inflammation (15), which could be attributed to the need of T_reg_ cells for the transition from M1 to M2 macrophage phenotype, the ability of T_reg_ cells to limit IFN-g and macrophage accrual, or the need for T_reg_ derived IL-10 (15–17). The regulation of myeloid and lymphoid immune cell infiltration and clearance works in concert with the stages of myogenesis for prompt muscle regeneration after acute injury.

By contrast, following large volume skeletal muscle loss that exceeds the regenerative threshold (volumetric muscle loss, VML), chronic inflammation produces an inhospitable microenvironment that does not allow for muscle regeneration. Instead, healthy muscle fibers are replaced by non-contractile fibrotic tissue resulting in chronic loss of function and permanent disability (18–20). This inflammatory microenvironment after VML has been previously characterized as having an enduring presence of CD4^+^ and CD8^+^ T cells, as well as improper macrophage polarization (6, 11, 15, 21, 22). Moreover, a muscle environment that is rich with both pro- and anti-inflammatory factors (i.e. TNF-α and TGF-β) can cause sustained presence of M2 macrophages that shift from ‘pro-reparative’ to ‘pro-fibrotic’ and drive FAPs to a pathological state rather than a regenerative one (23). However, the initial immune cell response following a critical muscle defect that often leads to chronic inflammation and fibrotic outcomes has not been elucidated. Previously, we characterized multiple muscle biopsy punch injuries in the mouse quadriceps as a model of VML. We found that a 2 mm injury caused damage to the tissue, but the muscle was able to regenerate without significant fibrotic scarring. However, a 3 mm injury caused persistent fibrotic scarring and inflammation through four weeks following injury (24).

A population of cells captured at the same time via high-dimensional immune cell profiling includes many distinct intermediate differentiation states of cells; however, classical analytical techniques only evaluate their average properties and thus mask trends occurring across individual cells and subpopulations (25–27). In this study, we approached this challenge by employing a unique dimensionality reduction and clustering analytical strategy that combines both uniform manifold approximation and projection (UMAP) and spanning-tree progression analysis of density-normalized events (SPADE); this analytical approach captures the functional heterogeneity of the early immune cell response and dynamics after an injury event and was applied to the scenario of critical VML injury (Figure S1). UMAP was utilized for graphing single cells isolated from multiparameter flow cytometric analysis into a lower dimensional representation, positioned by marker expression similarities, to visualize myeloid and lymphoid cellular densities and phenotype transitions over time in subcritical and critical VML (28). Furthermore, we leveraged SPADE to reconstruct cellular hierarchies and transitional states that can be inferred through time – a concept known as ‘pseudo-time’ (25, 29). By generating SPADE dendrograms with pooled sample data across timepoints, single cells were clustered into distinct nodes and then ordered along a trajectory to temporally track cellular lineages and progressions.

These dimensionality reduction and clustering tools provide greater insights into the underlying heterogeneous cell populations contributing to the early immune response after VML injury.

The objective of this study was to characterize the immune cell dynamics at early timepoints which led to the pathological phenotype following critical VML injury. We hypothesized that critical VML would lead to persistent elevation of key immune cell subpopulations, particularly those which are associated with promoting fibrosis.

## Results

### UMAP visualization of immune cell recruitment to critical VML injury reveals persistent myeloid cell response

Following subcritical (2 mm) or critical (3 mm) VML injury, immune cells were isolated from injured quadriceps muscles and quantified at 1, 3, and 7 days post injury via biplot gating of flow cytometry data. All CD11b^+^ myeloid cells from uninjured and injured animals at all timepoints (days 1, 3, and 7) were used to construct a UMAP plot that graphs single cells by their surface marker profiles, where further distances between cells indicates dissimilar cellular phenotypes (Figure 1). Surface marker expression values are represented on the UMAP, ranging from dark blue (low expression) to yellow (high expression), to visualize the position of myeloid cells with differing phenotypes throughout the plot (Figure 1A). Each CD11b^+^ cellular event is represented as a dot on the UMAP, with those coming from uninjured quadriceps overlaid in green and those from subcritical or critical VML injured quadriceps overlaid in blue and red, respectively (Figure 1B).

**Figure 1.**
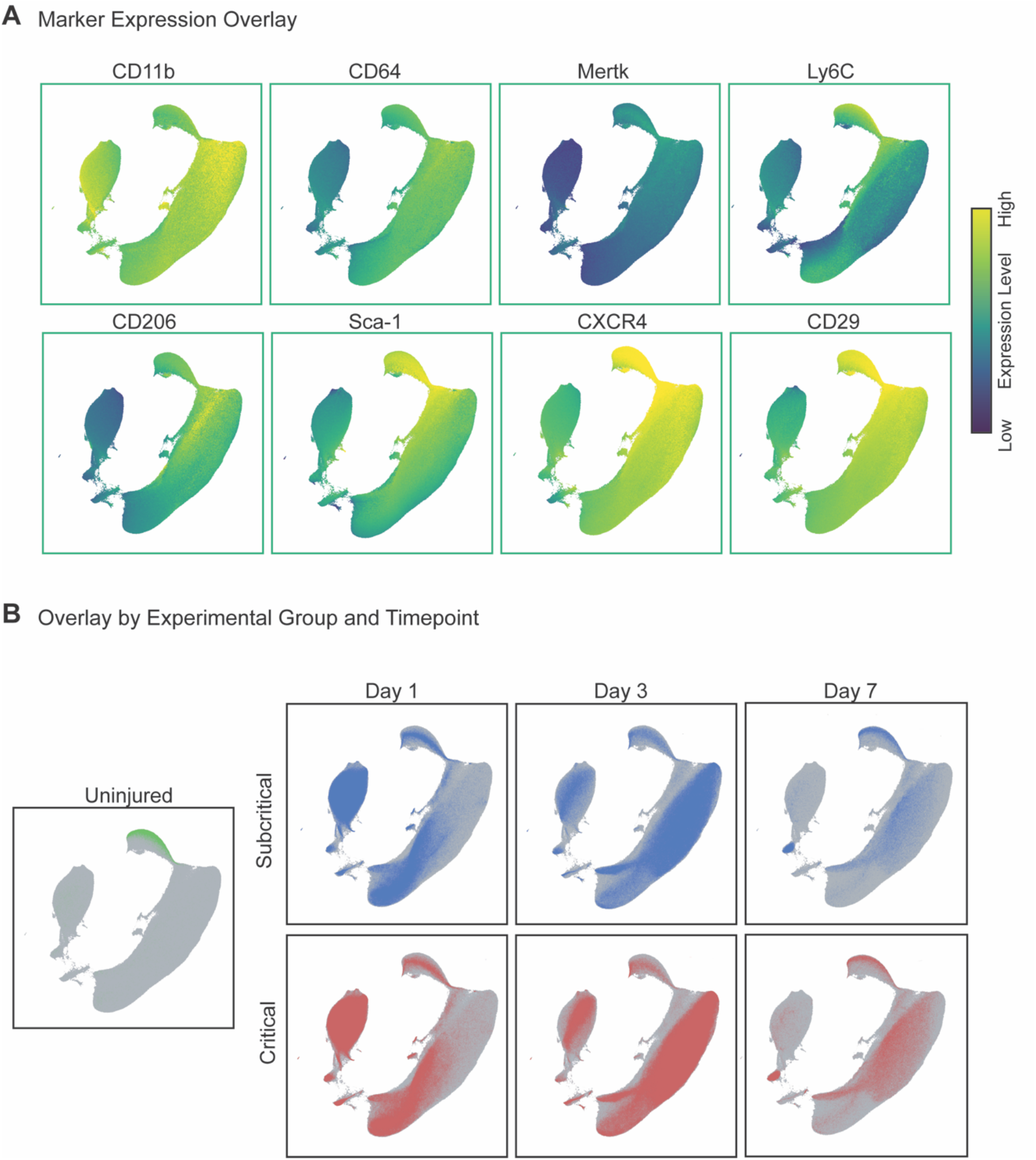
Persistent myeloid cell response in critical VML injury as visualized by UMAP analysis. UMAP representation comprised of single cell flow cytometry data from explanted quadriceps muscle at days 1, 3, and 7 post injury in addition to uninjured quadriceps. **(A)** UMAP projection overlaid with marker expression values of each targeted surface protein. Marker expression levels range from dark blue to yellow, representing low to high expression, respectively. **(B)** CD11b^+^ cells from uninjured quadriceps overlaid onto UMAP (green). CD11b^+^ cells from quadriceps that received a subcritical injury (blue, top) and those from quadriceps that received a critically sized VML injury (red, bottom) overlaid onto the UMAP at days 1, 3, and 7 post injury (left to right).

As visualized by UMAP, we observed that the cellular phenotypes of overall myeloid cells are consistent between injury sizes over time. This is indicated by the movement of events within the UMAP across timepoints, as myeloid cells from subcritical and critical VML injuries follow similar trajectories through UMAP space. At day 1, myeloid cells from injured quadriceps qualitatively localize near the bottom left of the UMAP plot, indicating that day 1 myeloid cells have low expression of CD206, MerTK, and CXCR4 but highly express Ly6C (Figure 1A). With time, the cells move towards the upper right portion of the UMAP, representing a phenotypic switch to increased CD206, CD64, MerTK, Sca-1, CD29, and CXCR4 expression and a lower expression of Ly6C by day 7. While the overall movement in the UMAP of CD11b^+^ events is similar between subcritical and critical injuries, the density of events is remarkably different. At days 3 and 7, critical VML injuries present with a higher density of myeloid cells within the UMAP compared to subcritical injuries, suggesting that inflammation has not been resolved by day 7.

The movement of CD11b^+^ cells as visualized by UMAP suggests an increase in mononuclear phagocyte populations persisting at days 3 and 7 as indicated by increased expression of CD64 (monocytes and macrophages) and MerTK (macrophages). Thus, a UMAP plot comprised of all CD11b^+^CD64^+^MerTK^−^SSC^lo^ monocytes and CD11b^+^CD64^+^MerTK^+^ macrophages pooled from all samples across all three timepoints was constructed (Figure 2). Expression levels of surface markers characterizing monocyte and macrophage subsets were overlaid onto UMAP plots to illustrate the location of unique mononuclear phagocyte subpopulations (Figure 2A). While the density of monocytes and macrophages appear similar between subcritical and critical VML injuries at days 1 and 3, there is a visibly increased density of these cells at day 7 in critical VML compared to subcritical (Figure 2B). This result suggests persistent inflammation in critical VML that is not observed in subcritical injuries. This retention of mononuclear phagocytes may be explained by the increased secretions of pro-inflammatory cytokines within the injury niche as indicated by an upwards trend in TNF-α expressing neutrophils at day 7 in critical VML (Figure S2). Notably, a subset of mononuclear phagocytes located at the far right of the UMAP is present in critical VML but not subcritical at day 7. This particular subset is characterized by low expression of Ly6C and high expression of CD206, suggesting an increase in non-classical monocytes and potentially pro-fibrotic M2-like macrophages induced by larger injury (Figure 2B).

**Figure 2.**
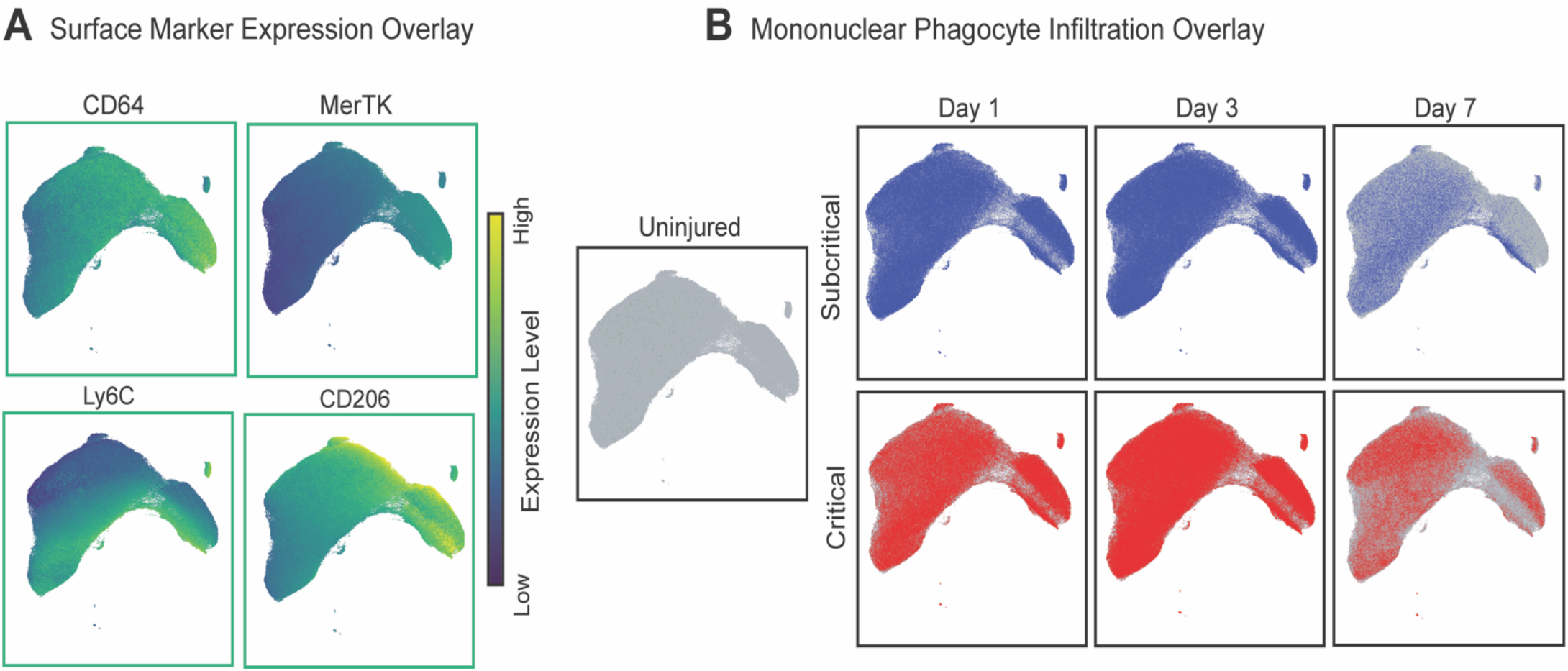
UMAP analysis reveals accumulation of mononuclear phagocytes in critically sized VML injuries. UMAP generated from flow cytometry gated CD11b^+^CD64^+^MerTK^+^ macrophages and CD11b^+^CD64^+^MerTK^−^SSC^lo^ monocytes extracted from quadriceps tissue (uninjured and at days 1, 3, and 7 post injury). **(A)** UMAP projection overlaid with surface maker expression values. Marker expression levels range from dark blue to yellow, representing low to high expression, respectively. **(B)** Visualization of mononuclear phagocyte infiltration in quadriceps that received no injury (uninjured), subcritical injury (blue, top) or critically sized injury (red, bottom) at each timepoint post injury (days 1, 3, and 7).

### Monocyte subpopulations facilitate the chronic inflammation characteristic of critical VML

To overcome the intrinsic challenge of subjective cell subset identification presented by traditional methods for analyzing multiparameter single-cell data, unsupervised SPADE analysis was applied to our flow cytometry data. SPADE groups the cells into distinct clusters, or nodes, by similar marker expression and then arranges these nodes into branching trajectories that infer cellular transitions through time (30). We constructed a SPADE dendrogram with CD11b^+^CD64^+^MerTK^−^ SSC^lo^ monocytes from all samples and all timepoints (days 1, 3, and 7) after injury, including uninjured controls. The SPADE dendrogram is color annotated into three monocyte subpopulations identified by the Ly6C expression of each node, as shown in the corresponding heatmap where each column correlates to one SPADE node (Figure 3A, B). From left to right, the phenotype of the monocyte SPADE nodes transition from Ly6C^hi^ to Ly6C^lo^, illustrating the dynamic biological progression of monocytes after VML.

**Figure 3.**
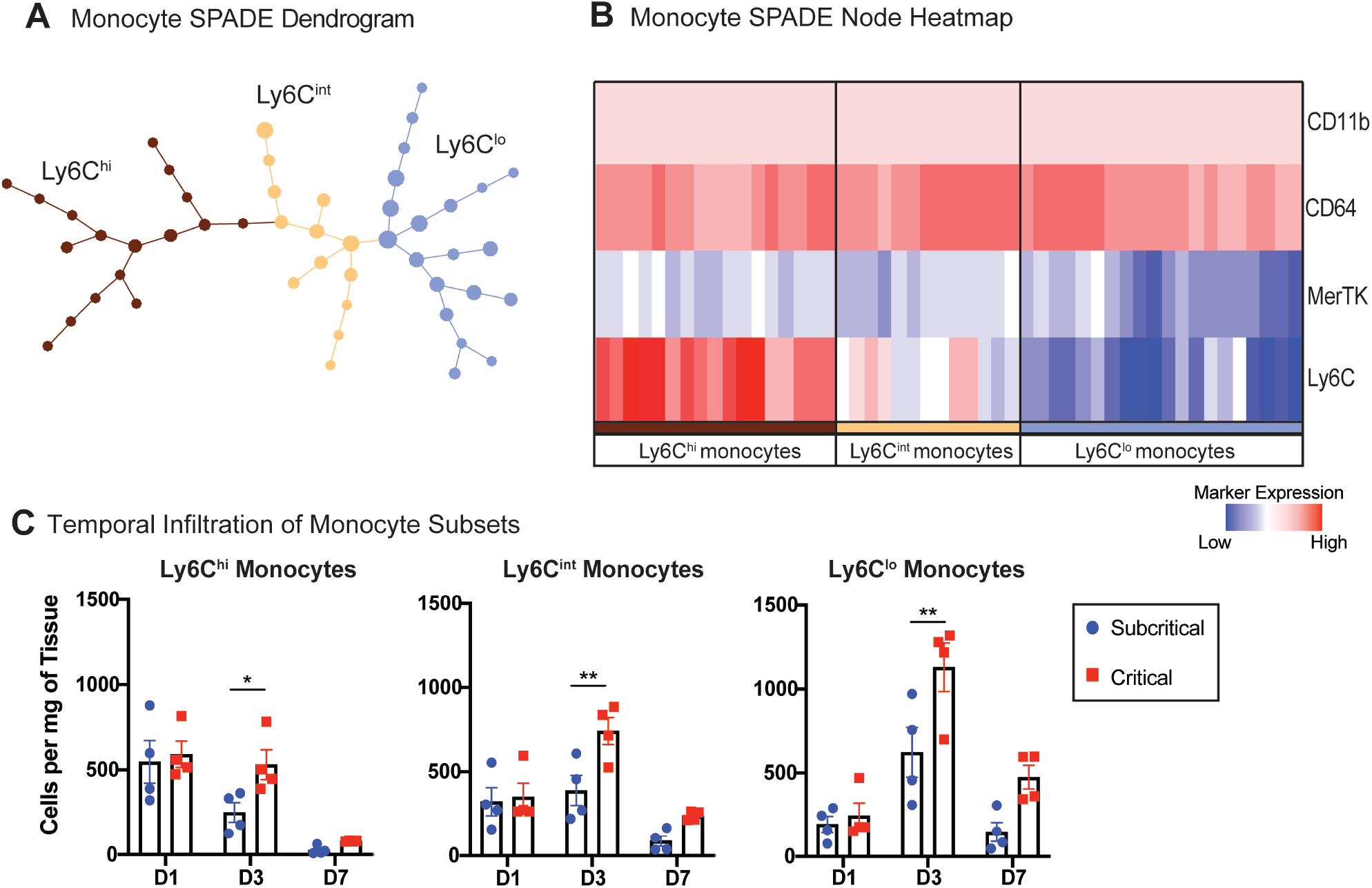
Unbiased SPADE clustering identifies distinct monocyte subpopulations that are significantly increased in critically sized injuries 3 days post VML. SPADE dendrogram generated from all CD11b^+^CD64^+^MerTK^−^SSC^lo^ monocytes isolated from quadriceps muscle of mice which were uninjured or those which received a subcritical or critically sized injury (at days 1, 3, and 7 post injury). **(A)** Monocyte SPADE dendrogram annotated by nodes containing monocytes with low (Ly6C^lo^), intermediate (Ly6C^int^), or high (Ly6C^hi^) expression of Ly6C. **(B)** Protein signature of each SPADE node represented as a heatmap. SPADE nodes classified according to their expression of Ly6C. Color annotation of SPADE nodes on the dendrogram align with those on the heatmap. **(C)** Quantification of monocyte subsets (per mg of tissue) within quadriceps muscle at each timepoint post injury. Data presented as mean ± S.E.M. Statistical analyses performed include two-way ANOVA with *Sidak* multiple comparisons between injury sizes at each timepoint. \**p*<0.05, \*\**p*<0.01. n=4 animals per experimental group. D1: day 1, D3: day 3, D7: day 7.

The concentration of cells within each of the three SPADE-identified monocyte subpopulations was quantified for all timepoints (Figure 3C). For all subsets, there was a significant increase in the number of monocytes present in critical VML at day 3 post injury. Ly6C^int^ monocytes – which would typically be left out of traditional gating strategies between the ‘high’ and ‘low’ expressing gates – and Ly6C^lo^ monocytes both reached peak concentrations at day 3 with Ly6C^lo^ monocytes reaching the highest concentration of all the subpopulations at any timepoint. To evaluate the cytokine expression profile of these monocyte subsets, we again performed flow cytometry analysis from explanted quadriceps at days 3 and 7 after subcritical or critical VML. There were no differences found in the concentration of TNF-α^+^ or SDF-1^+^ monocyte subsets infiltrating subcritical and critical VML injuries at day 3 (Figure S3). However, at day 7, the number of Ly6C^int^ monocytes expressing TNF-α and Ly6C^hi^ monocytes expressing SDF-1 were significantly elevated in critical VML. Almost no monocytes, regardless of subset classification, expressed TNF-α or SDF-1 in subcritical injuries at day 7 (Figure S3).

### Pseudo-time analysis identifies distinct macrophage subsets dysregulated following critical VML injury

All CD11b^+^CD64^+^MerTK^+^ macrophages, from uninjured and injured quadriceps (at days 1, 3, and 7 post injury) were used to generate a SPADE dendrogram (Figure 4A). SPADE clustered the macrophages into nodes ordered along 4 marked trajectories, each color-coded, characterized by the surface marker expression of each node (Figure S4). Most macrophages present within the injured muscle at day 1 are clustered within nodes of the initial M1-like trajectory, characterized by its Ly6C^hi^CD206^lo^ expression profile. Moving down the dendrogram, as indicated by dotted gray arrows, the ordered nodes split into 3 separate branches: an unpolarized (Ly6C^lo^CD206^lo^) trajectory, an M2-like (Ly6C^lo^CD206^hi^) trajectory, and a ‘hybrid’ (Ly6C^hi^CD206^hi^) macrophage trajectory (Figure 4A). Identification of both unpolarized and hybrid macrophage phenotypes is often overlooked with traditional, manual bi-plot gating strategies applied to flow cytometry data; with SPADE clustering and pseudo-time trajectory visualization, these heterogeneous subtypes are more readily discovered. Quantifying the number of macrophages within each annotated SPADE trajectory, it was found that there were no changes in the concentrations of M1-like or unpolarized macrophages between injury sizes at any timepoint. By contrast, both M2-like macrophages and hybrid macrophages were significantly elevated at 7 days post injury (Figure 4B). Our data shows that CXCR4 expression elevates concurrently with CD206 for both M2-like and hybrid macrophage subtypes (Figure S4). Thus, the dual expression of CD206 and CXCR4 may define specific macrophage subsets with a sustained and dysregulated response following critical VML.

**Figure 4.**
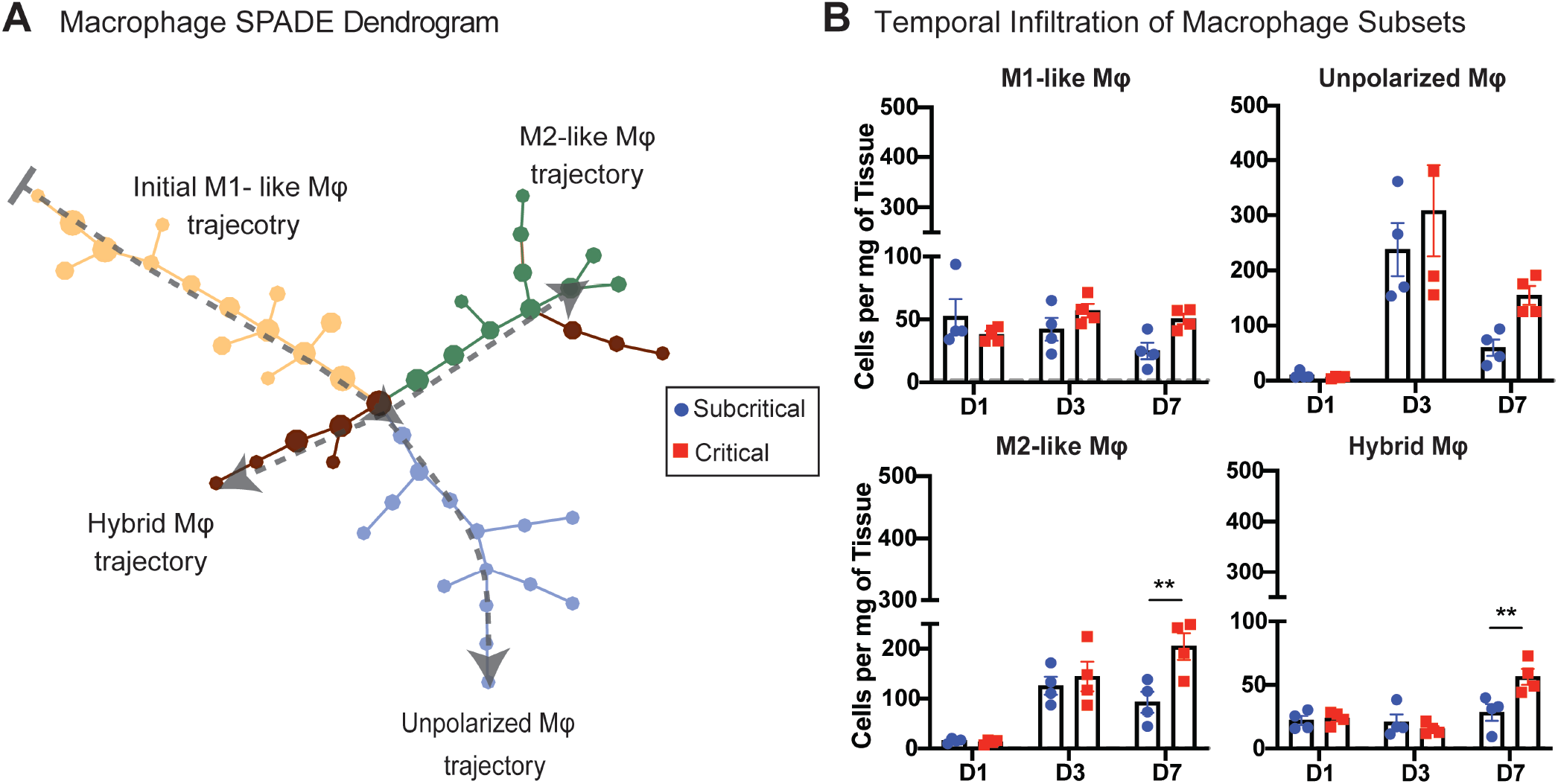
SPADE-identification of macrophage subsets uncovers two unique populations expressing CD206 which are significantly increased in critically sized injuries at day 7. SPADE dendrogram generated from all CD11b^+^CD64^+^MerTK^+^ macrophages isolated from quadriceps muscle of mice which were uninjured or those which received a subcritical or critically sized injury (at days 1, 3, and 7 post injury). **(A)** Macrophage SPADE dendrogram annotated based on CD206 and Ly6C expression to designate nodes to a M1-like, M2-like, unpolarized, or hybrid macrophage phenotype. Gray arrow indicates general movement over time following injury. **(B)** Quantification of macrophage subsets (per mg of tissue) within quadriceps muscle at each timepoint post injury. Data presented as mean ± S.E.M. Statistical analyses performed include two-way ANOVA with *Sidak* test for multiple comparisons between injury sizes at each timepoint. \*\**p*<0.01. n=4 animals per experimental group. D1: day 1, D3: day 3, D7: day 7, Mφ: macrophage.

While most studies have characterized the role of M2-like macrophages in establishing an antiinflammatory microenvironment and promotion of tissue regeneration, in the presence of chronic inflammatory stimuli, M2-like macrophages are known to secrete large amounts of pro-fibrotic cytokines and promote tissue and organ fibrosis (31). To examine the intracellular cytokine profile of the significantly elevated M2-like macrophages in critical VML, flow cytometry analysis was performed at days 3 and 7 post subcritical and critical VML injury. A SPADE dendrogram was constructed of all CD11b^+^CD64^+^MerTK^+^ macrophage events from both timepoints and injury sizes, and nodes were grouped into the same four phenotype subtypes (M1-like, M2-like, unpolarized, and hybrid) based on expression of Ly6C and CD206 (Figure 5A).

**Figure 5.**
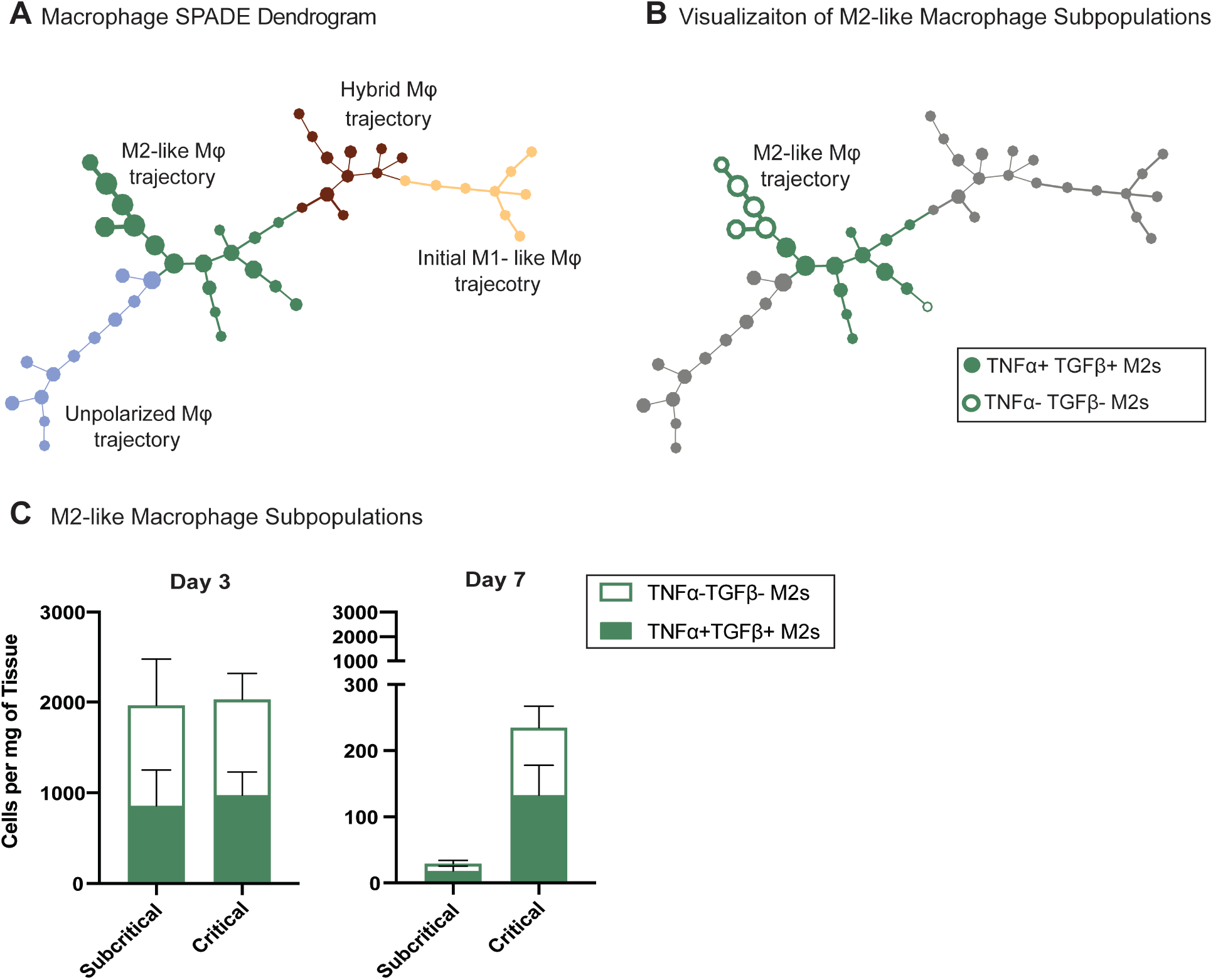
Accumulation of TNF-α^+^ and TGF-β^+^ M2-like macrophages may promote pathological fibrosis in critical VML defects. Flow cytometry was performed on single cells isolated from quadriceps of mice that received a subcritical or critically sized VML injury at days 3 and 7 post injury. CD11b^+^CD64^+^MerTK^+^ macrophages pooled from all samples from both injury sizes and both timepoints were used to construct a SPADE dendrogram. **(A)** Macrophage SPADE dendrogram annotated based on CD206 and Ly6C expression to designate nodes to a M1-like, M2-like, unpolarized, or hybrid macrophage phenotype. **(B)** M2-like macrophages (green annotation) distinguished by their intracellular expression of TNF-α and TGF-β (TNF-α^+^ TGF-β^+^ nodes: solid green; TNF-α^−^ TGF-β^−^ nodes: green outline). **(C)** Number of TNF-α^+^ TGF-β^+^ and TNF-α^−^ TGF-β^−^ M2-like macrophages per mg of tissue represented as a stacked bar graph at day 3 and day 7 post subcritical or critical VML injury. Data presented as mean ± S.E.M. n= 3-4 animals per experimental group. Mφ: macrophage.

Within the M2-like macrophage subset, nodes were further annotated based on expression of intracellular TNF-α and TGF-β. M2-like macrophage SPADE nodes that expressed both TNF-α and TGF-β were annotated as solid green whereas nodes that did not express either cytokine were annotated with only a green outline (Figure 5B). All nodes either expressed both or neither of the two cytokines, as there were no TNF-α^+^TGF-β^−^ or TNF-α^−^TGF-β^+^ expressing M2-like nodes. At day 3 post injury, there were no differences in the concentrations of TNF-α^+^TGF-β^+^ or TNF-α^−^ TGF-β^−^ M2-like macrophages between injury sizes. At day 7, very few M2-like macrophages remain in subcritical defects, but there is a clear retention of M2-like macrophages in critical VML injured quadriceps-the majority of them expressing both TNF-α and TGF-β (represented as green segment of stacked bar graph) (Figure 5C).

To further evaluate the sustained CD206^+^ cellular infiltration into critical VML injuries, subcritical and critical VML injured quadriceps cross-sections were used for immunohistochemical (IHC) analysis at 7 days post injury (Figure 6). Cross-sections were stained with phalloidin (gray), DAPI (blue), and CD206 (red). Uninjured control quadriceps showed healthy skeletal muscle morphology with DAPI^+^ myonuclei located at the periphery and little to no CD206^+^ cells (Figure 6A). Quadriceps that received subcritical injuries presented with visible DAPI^+^ cellular infiltration between myofibers at day 7. Some of these cells, present within the defect space or between surrounding myofibers, also expressed CD206^+^. The morphology of myofibers adjacent to the defect space appear largely unaffected. (Figure 6B). In contrast, there was a qualitative increase in the infiltration and persistence of CD206^+^ cells within the defect area of critically injured quadriceps. There is an obvious lack of tissue structure with myofibers spaced apart, distorted in their morphology, and with clusters of mononuclear cells surrounding them (Figure 6C). Taken together, with critical VML injury, we observed an increase in CD206^+^ cell retention within the injury niche at day 7, quantitatively through SPADE pseudo-time analysis and qualitatively through IHC.

**Figure 6.**
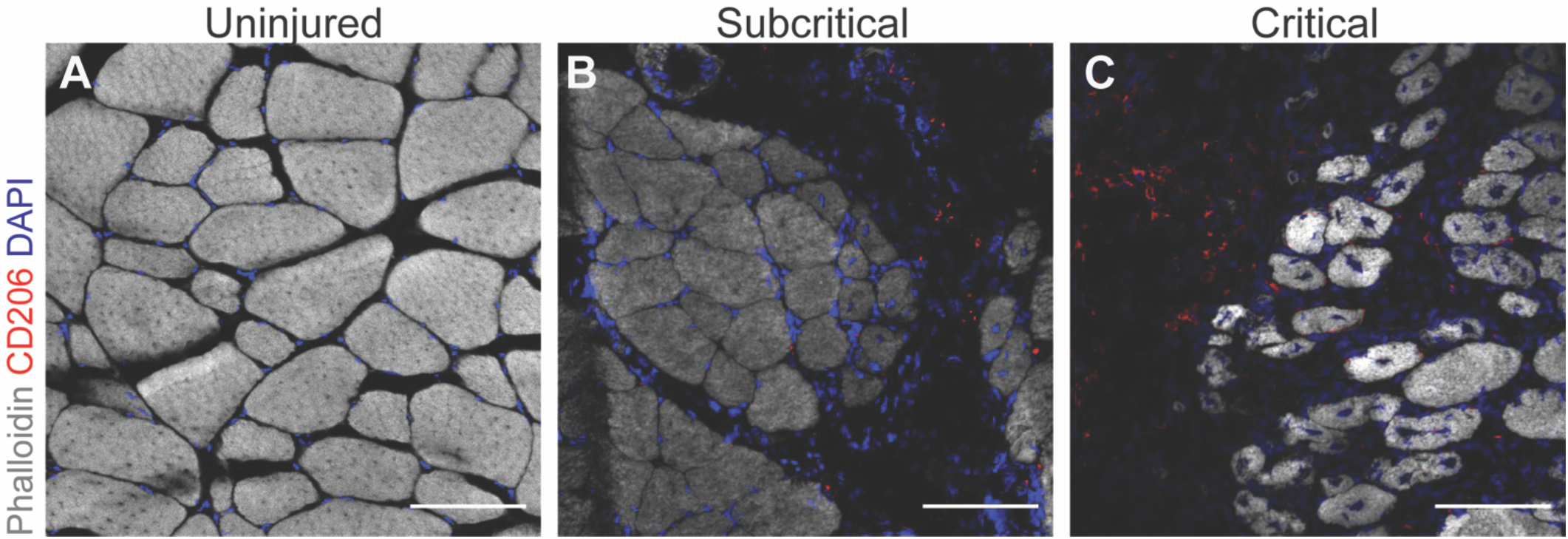
Sustained CD206^+^ cellular infiltration to injury milieu of critically sized defects. Representative images of quadriceps cross-sections from uninjured control **(A)**, subcritical injury **(B)**, and critically sized injury **(C)** at a day 7 timepoint. Cross-sections stained with phalloidin (gray), CD206 (red), and DAPI (blue). Scale bars represent 100 μm.

### Chronic inflammatory stimuli from critical VML propagates unchecked T cell activation and recruitment to injury milieu

All CD3^+^ T cells from all animals and at all timepoints (days 1, 3, and 7) were used to construct a UMAP plot to visualize T cell infiltration and phenotype transitions in a lower dimensional space (Figure 7). Relative expression levels of each measured T cell surface marker were overlaid onto the UMAP to illustrate the locations of heterogeneous T cell subpopulations within the UMAP space (Figure 7A). The few T cells present in uninjured quadriceps were overlaid in green on the UMAP while T cells from subcritical injured quadriceps are overlaid in blue, and T cells from critical injured quadriceps are overlaid in red, by timepoint (Figure 7B). The density and location of overlaid T cells from different biological samples provides a visual representation of T cell subtype dynamics following subcritical and critical injury.

**Figure 7.**
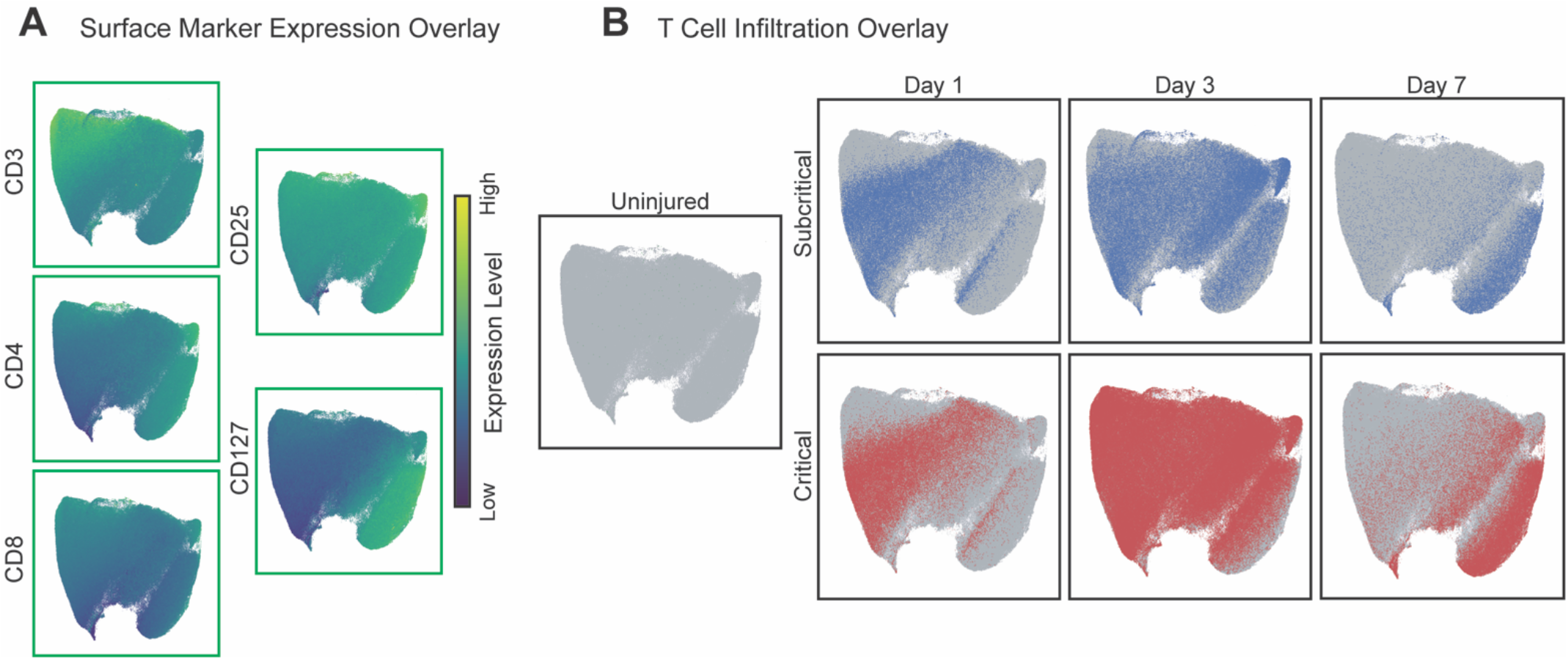
Critical VML injury presents with increased T cell recruitment 3 days post injury as visualized by UMAP. CD3^+^ T cells pooled from all samples (uninjured, subcritical, and critical VML injured quadriceps) and all timepoints (days 1, 3, and 7) were used to generate a UMAP projection. **(A)** Expression levels of T cell surface markers overlaid onto UMAP. Marker expression levels range from dark blue to yellow, representing low to high expression, respectively. **(B)** CD3^+^ T cells isolated from uninjured quadriceps (green), subcritical injuries (blue, top), and critically sized injuries (red, bottom) at each timepoint.

Temporal dynamics of T cells were observed through the UMAP, as the T cells move from left to right through the UMAP space over time (Figure 7B). T cells from subcritical and critical VML share similar surface marker dynamics, indicated by their similar location within the UMAP at each timepoint. At day 1, it appears that T cells have a low expression of CD4 and CD127. Yet by day 3, there is a clear shift in phenotype as T cells increased their expression of CD4 and CD127, with CD25 and CD8 remaining relatively constant with time (Figure 7A, B). Importantly, there was a visible increase in T cell density in critical VML at days 3 and 7 post injury as observed in the UMAPs (Figure 7B) which may indicate hyperactivation of the adaptive immune response.

To characterize and quantify unique subsets of T cells within subcritical and critical VML injuries, all CD3^+^ T cells from all animals and timepoints were used to generate a SPADE dendrogram (Figure 8A). SPADE clustering of T cell events by their surface marker expression profile (Figure 8B) also revealed a clear localization of T cells by timepoint indicating the phenotypic transition that occurs within the first week following injury. The nodes were grouped into 3 time-associated clusters: the initial T cell response (annotated red), transition T cells (annotated green), and final T cell fates (annotated blue) (Figure 8A). The percentage of total T cells within each of these time-associated clusters was quantified (Figure 8C). The percentage of total T cells from day 1 samples in the initial T cell response population was significantly higher than the percentage of T cells from 3 and 7 days post injury, and the percentage of day 7 T cells is significantly lower than those at day 3 (Figure 8C). This result verifies that the left side of the SPADE dendrogram (red annotation) is comprised predominately of day 1 T cells. Within the transitional T cell population, representing a group of T cells that are not yet in their ‘final’ phenotype state, the percentage of T cells from days 3 and 7 were significantly higher than those from day 1 (Figure 8C). In final T cell fate nodes (far right of dendrogram), the percentage of T cells at day 7 were significantly elevated compared to the percentage at days 1 and 3 (Figure 8C). These findings confirm that the clustering of T cells via surface marker expression coincides with the progression of time (indicated with dotted gray arrow). No differences were observed in the percentage of T cells between subcritical and critical injuries at any time point within these time-associated node clusters.

**Figure 8.**
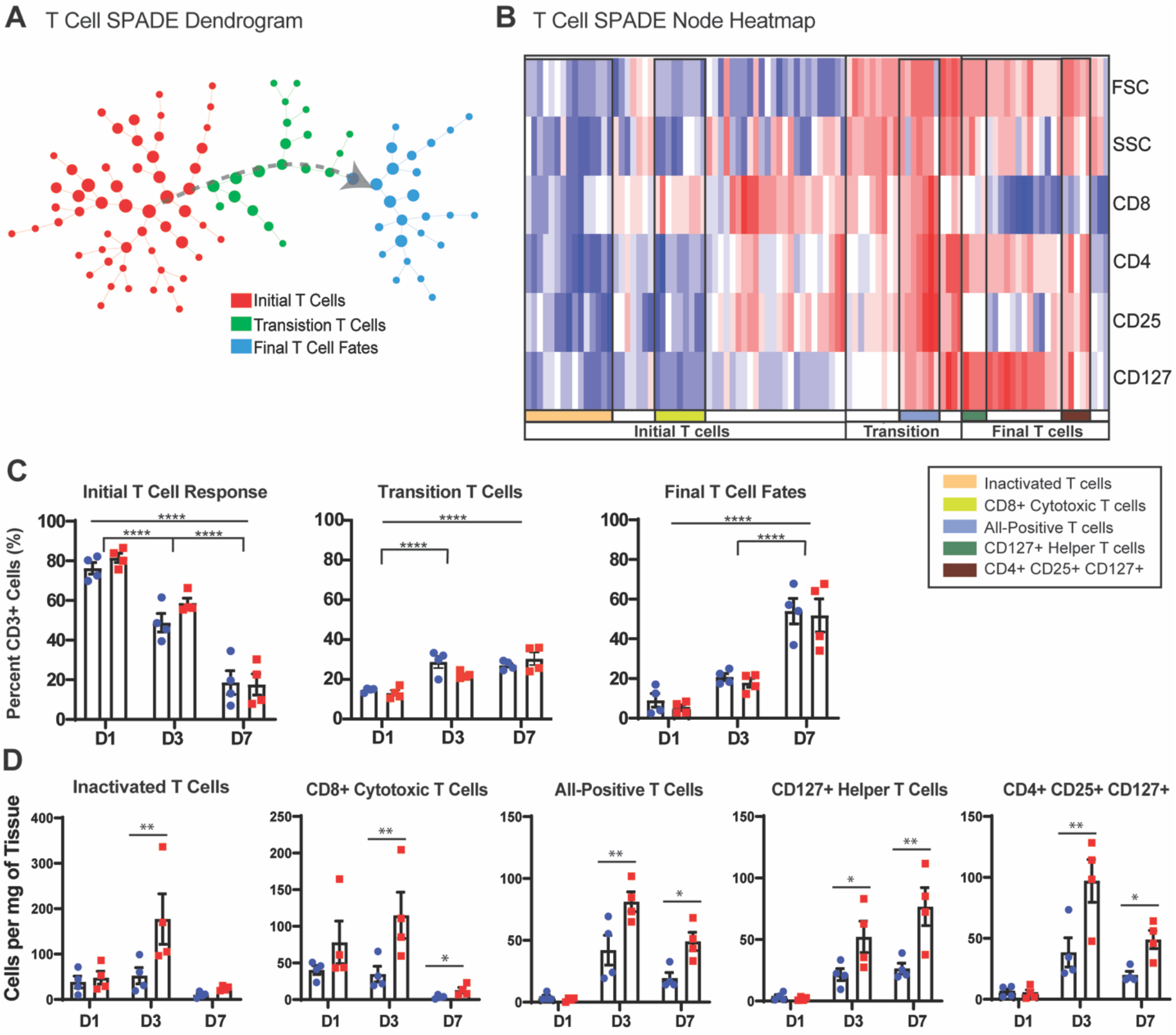
Pseudo-time analysis reveals a dysregulated T cell response to critical VML injury. Pre-gated CD3^+^ T cells from all samples (uninjured, subcritical, and critical injuries) and all timepoints (days 1, 3, and 7) were used to construct a SPADE dendrogram. (A) SPADE nodes annotated by the relative percentage of T cells present from each timepoint. Gray arrow indicates general movement over time following injury. (B) Protein signature of each SPADE node represented as a heatmap. SPADE nodes grouped by temporal infiltration kinetics (initial T cells, transition T cells, or final T cell fates) and further annotated by marker expression profiles. (C) The percentage of total T cells from each timepoint within each annotated region of the SPADE dendrogram. (D) T cell subtype cell counts quantified as cell concentration (cells/mg tissue). Quantified subtypes identified and annotated in B. Data presented as mean ± S.E.M. Statistical analyses performed as two-way ANOVA with *Sidak* multiple comparisons test to determine differences between injury sizes. \**p*<0.05, \*\**p*<0.01, \*\*\*\**p*<0.0001, n=4 animals per experimental group. D1: day 1, D3: day 3, D7: day 7.

Within each of the three time-associated node clusters, specific T cell subtypes of interest were identified by their unique marker expression profile as shown at each SPADE node in the form of a heatmap (Figure 8B). The SPADE nodes of the heatmap are organized from left to right in the same progression as the dendrogram. Within the greater ‘initial T cell response’ grouping, inactivated T cells were characterized by their low expression of all measured markers in addition to low forward scatter area (FSC) and side scatter area (SSC), which indicate small relative cell size and low intracellular granularity, respectively. The cells present in these nodes were quantified and it was found that there were significantly more inactivated T cells in critical VML at 3 days post injury compared to subcritical injuries (Figure 8D). Nodes containing CD8^+^ cytotoxic T cells were identified by their high expression of CD8 but low expression of all other surface markers, FSC, and SSC (Figure 8B). CD8^+^ cytotoxic T cells were significantly elevated at both days 3 and 7 following critical VML injury (Figure 8D).

As the dendrogram progresses into transition and final T cell fate stages, it is observed that cell size and intracellular granularity is increased as measured by FSC and SSC (Figure 8B). A subpopulation of transitional T cells positive for all measured surface markers was identified. These ‘all-positive T cells’ were present within critically injured muscle at significantly higher concentrations at days 3 and 7 (Figure 8B, D). Two subsets of interest were identified within the final fate T cells comprised chiefly of T cells from day 7. One of these subsets highly expressed both CD4 and CD127 (Figure 8B, green annotation). These CD127+ helper T cells were significantly increased at days 3 and 7 after critical VML (Figure 8D). Finally, a group of nodes containing CD127^+^ T_reg_ cells (CD4^+^CD25^+^) were identified within the greater final fates T cells clustering. This T_reg_ subset presented with low expression of CD8 and was also significantly elevated at days 3 and 7 post injury in critical VML (Figure 8B, D). Taken together, SPADE analysis was able to reconstruct the phenotypic transitions of T cell populations occurring through time and facilitated robust characterization and quantification of specific T cell subsets to reveal a dysregulated T cell response occurring at day 3 and 7 in critical VML.

## Discussion

The proper coordination of immune cells is crucial for prompt and proper regeneration and repair of damaged muscle following minor muscle injuries. Although skeletal muscle possesses remarkable regenerative capabilities, the traumatic loss of muscle characteristic of VML ablates the extracellular matrix and MuSC niche necessary for the initiation of myogenesis; thus, muscle’s innate capacity for regeneration becomes inadequate to functionally recover muscle (32). In this series of studies, our data reveals key differences in the concentrations and temporal dynamics of identified immune cell subsets present within injured muscle following subcritical and critical VML injury. Subcritical injuries elicited an immune response similar to what is expected from an acute muscle injury, as most of the inflammation had been resolved by day 7. In contrast, critical VML presented with a sustained elevation of several unique myeloid and lymphoid cell subsets as characterized by UMAP visualization and SPADE pseudo-time analysis. These results were summarized in Figure S5, where the average concentration data for each subtype and injury size were plotted to demonstrate the altered immune response to critical VML.

While single cell resolution by flow cytometry analysis provides a powerful tool for temporal evaluation of the immune response, increased dimensionality obscures the underlying heterogeneity of the data and increases the likelihood of introducing user bias when identifying cell populations (33). We harnessed the unbiased dimensionality reduction methods, UMAP and SPADE, to analyze the temporal progression of immune cell frequencies and phenotypic transitions present in injured muscle dependent on injury size. Visualizing immune cell heterogeneity as a lower dimensional representation by UMAP, we observed that critical VML injury showed increased CD11b^+^ cell retention in injured muscle at day 7 and a transient increase in MerTK, CXCR4 and CD206 expression (Figure 1). We found that in particular, mononuclear phagocytes elicited a prolonged immune response in critical VML, including a specific CD206^hi^Ly6C^lo^ subset that was not present in subcritical injuries by day 7 (Figure 2), a finding that would not be easily discovered without implementation of advanced, dimensionality reduction visualization techniques.

We hypothesized that the sustained presence of mononuclear phagocytes within critical VML injuries may lead to the persistence of pro-inflammatory cytokines such as TNF-α and SDF-1. There was a significant increase in all identified monocyte subsets at day 3 post injury, suggesting an increased extravasation of monocytes from circulation to larger injuries (Figure 3C). We found that at day 7 post injury, the concentrations of TNF-α^+^ neutrophils in addition to TNF-α^+^ and SDF-1^+^ monocytes were significantly elevated in critical VML but little to none were present in subcritical injuries (Figure S2, S3). These results may indicate that early accumulation of pro-inflammatory myeloid cells in critical VML propagates the secretions of potent leukocyte chemoattractants such as TNF-α and SDF-1, leading to aberrant immune cell retention at day 7 onwards. In an environment rich in cytokines such as TNF-α and IFN-g, it is likely that MuSCs may efficiently activate and proliferate but fail to differentiate into myotubes as a result of a failed inflammatory-to-regenerative transition (1, 4); thus, myogenesis is impaired and fibrotic and fatty infiltration fill the defect rather than functional muscle.

In minor injuries, elevation of M2-like macrophages and their progenitors, non-classical Ly6C^lo^ monocytes, has been linked with improved muscle healing (10, 34). However, our results indicate an abnormal persistence of M2-like macrophages in critical VML at day 7 (Figure 4, 6). Taking advantage of the unique capability of SPADE to preserve rare cell types often masked in bulk cellular analysis, we discovered a population of hybrid CD206 and Ly6C co-expressing macrophages which were also significantly increased in critical VML (Figure 4). We found that both of these CD206^hi^ macrophage subsets have concurrent elevations in CXCR4 expression (Figure S4) which has been reported to be linked to fibrosis, in part through their expression and secretion of tissue inhibitor of metalloprotease 1 (TIMP1) (35). It is notable that CXCR4 is elevated in M2s, as they are known to secrete pro-fibrotic cytokines such as TGF-β. It is also of interest that CXCR4 is highly expressed in hybrid macrophages, suggesting a potential role for CD206^hi^Ly6C^hi^ macrophages in fibrosis.

Previous studies have shown that macrophages simultaneously expressing pro- and antiinflammatory cytokines may result from impaired M1-to-M2 phenotypic transitions and contribute to chronic inflammation and subsequent tissue fibrosis (6, 36, 37). We sought to further characterize the cytokine profile of the M2-like macrophages persisting in critical VML and found that the majority of M2-like macrophages present at day 7 co-expressed TNF-α and TGF-β (Figure 5). It has been shown that when macrophages co-express these cytokines, TGF-β dominates the response and leads to unregulated ECM deposition by FAPs to facilitate tissue fibrosis (6). Moreover, it has been shown that the chronic activation of M2-like macrophages can activate muscle-resident fibroblasts via their release of TGF-β (38) which may further exacerbate fibrosis and a failed regenerative outcome. Future studies probing the cytokine signaling between FAPs and persistent macrophage subsets are necessary to elucidate the role of M2-like and hybrid macrophages in critical VML.

T cells are known to play important roles in aiding macrophage trafficking and polarization during muscle healing (14, 17). Further, as phagocytic macrophages populate the injury, they present antigens on their cell surface and secrete cytokines responsible for activating T cells and the adaptive immune response (39). We found that overall T cell numbers peaked around day 3 post injury, as would be expected (14). Despite seemingly appropriate dynamics, we found that there were several subsets of T cells that presented with an altered response to critical VML at days 3 and 7 compared to subcritical injuries (Figure 7, 8). We discovered that the surface expression characterizing T cell subpopulations changed as a function of time, as represented by the movement of T cells through the SPADE dendrogram from left to right (Figure 8). This lineage of inferred cellular transitions reconstructed by SPADE is known as pseudo-time and can be utilized to assess cellular progressions although data was collected at snapshots in time (25).

We qualitatively observed an increase in T cell size over time, represented by FSC, which indicates antigenic stimulation and T cell activation (40). Both CD8^+^ cytotoxic T cells and a population of T cells highly expressing all measured markers were significantly elevated in critical VML at days 3 and 7 (Figure 8), indicating increased T cell activation induced by larger injury size. CD4^+^CD8^+^ T cells have been shown to be highly cytotoxic which may lead to intensified tissue damage (41), but further investigation is necessary to determine if these double-positive T cells are detrimental to muscle regeneration. Lastly, we identified two T cell populations expressing IL-7 receptor, CD127: CD127^+^CD4^+^ helper T cells and CD127^+^CD4^+^CD25^+^ T_reg_ cells (Figure 8). As CD127 has been reported to be expressed on activated T_reg_ cells in the presence of IL-7 (42), our results may be indicative of elevated IL-7 in our critical VML model. IL-7 has been found to reduce myoblast differentiation and fusion (43), so future studies measuring whether increased IL-7 induced by critical VML upregulates CD127^+^ T cells and impedes myogenesis is of interest. While significant increases in T_reg_ cells after critical VML was not expected, it is possible that despite a local enrichment of T_reg_ cells, their immunosuppressive function is reduced in the presence of chronic inflammatory stimuli (44).

We have previously reported that 3 mm full thickness defects in the murine quadriceps VML model results in fibrosis, fatty infiltration, and lack of myofiber regeneration within the injury site-similar to the clinical scenario (24). Here, we elucidate the key immune cellular players that underlie this pathophysiology. Uncovering specific immune cell subtypes dysregulated in critical VML, particularly those with heterogeneous phenotypes overlooked with traditional gating strategies, is a novel approach to examining failed endogenous repair mechanisms at the cellular and molecular level. These studies may provide the necessary foundation for the development of targeted regenerative immunotherapies to improve clinical outcomes following VML.

## Methods and Materials

### Animals

C57BL/6J mice were purchased from Jackson Laboratories and maintained as a breeding colony. All animals used in the study were male, 6.1±0.5 (mean ± standard deviation) months in age at the time of euthanasia. All animals were used according to the protocols approved by the Georgia Institute of Technology Institutional Animal Care and Use Committee.

### Quadriceps volumetric muscle loss injury

Surgical procedure performed as previously reported (24). Briefly, the left hindlimb was prepped and sterilized. A single incision was made above the quadriceps and a 2 mm or 3 mm biopsy punch (VWR, 21909-132, −136) was used to make a full-thickness muscle defect. Skin was closed with wound clips and muscle was left to recover without intervention for 1, 3, or 7 days before euthanasia by CO_2_ inhalation.

### Tissue harvest andflow cytometry

Samples were prepared for flow cytometry analysis on a FACS AriaIII flow cytometer (BD Biosciences) as previously reported (45). Briefly, entire injured (or uninjured for controls), left quadriceps were harvested and digested with 5,500U/ml collagenase II and 2.5U/ml Dispase II for 1.5 hours in a shaking 37°C water bath. The digested muscles were filtered through a cell strainer to obtain a single cell suspension. Single-cell suspensions were stained for live cells using Zombie NIR (BioLegend) dyes in cell-culture grade PBS per manufacturer instructions. Cells were then fixed in 4% PFA for 10 minutes at 4°C. Cells were stained with cell phenotyping antibodies in a 1:1 volume ratio of 3% FBS and Brilliant Stain Buffer (BD Biosciences) according to standard procedures. The following antibodies were used for T cell phenotyping: BV605-conjugated anti-CD4 (BioLegend), BV785-conjugated anti-CD8 (BioLegend), BV421-conjugated anti-CD3ε (BioLegend), PerCP-Cy5.5-conjugated anti-CD25 (BioLegend), and APC-conjugated anti-CD127 (BioLegend). The following antibodies were used for myeloid cell phenotyping: BV421 or PE-Cy5-conjugated anti-CD11b (BioLegend), APC-Cy7-conjugated anti-Ly6G, BV510 or PerCP-Cy5.5-conjugated anti-Ly6C (BioLegend), BV711 or FITC-conjugated anti-CD64 (BioLegend), PE or APC-conjugated anti-MerTK (BioLegend), PE-Cy7 conjugated anti-CD206 (BioLegend), FITC-conjugated anti-Ly6A/E (BioLegend), APC-conjugated Lineage antibody cocktail (BD Pharmigen), APC-conjugated anti-CD31 (BioLegend), PE-Cy5 conjugated anti-CD29 (BioLegend), and PerCP-Cy5.5-conjugated anti-CXCR4 (BioLegend). The following intracellular antibodies were used in flow cytometry experiments when indicated: BV510-conjugated anti-TNF-α (BioLegend), BV421-conjugated anti-TGF-β (BioLegend), and PE-conjugated anti-SDF-1 (R&D Systems). 30μL of CountBright Absolute Counting Beads (C36950, Invitrogen) were added per sample for absolute quantification of cell populations.

### Immunophenotyping of myeloid and lymphoid cell subsets

Single, live cells were selected in FlowJo software for subsequent immunophenotyping analysis. Myeloid cells were identified as CD11b^+^ cells while lymphoid populations were identified as CD3^+^. Neutrophils were selected as CD11b^+^Ly6G^+^ cells. Monocytes were gated as CD11b^+^CD64^+^MerTK^−^SSC^lo^ cells and macrophages as CD11b^+^CD64^+^MerTK^+^ cells. Subsets of monocytes and macrophages were further analyzed using SPADE analysis, as described.

### Uniform Manifold Approximation and Projection (UMAP)

UMAPs generated as previously reported (27, 45, 46). Briefly, UMAP was used to embed high-dimensional flow cytometry data into a space of two dimensions, cells are visualized in a scatter plot where similarity is demonstrated via proximity to other points. Prior to UMAP dimensional reduction, each flow cytometry sample was pre-gated to select cellular subsets of interest (i.e. CD11b^+^ myeloid cells and CD3^+^ T cells) and then imported into Python 3.7 using fcsparser (https://github.com/eyurtsev/fcsparser) and Pandas 2.5. Each channel except for FSC and SSC was normalized by applying arcsinh/150. UMAP parameters of n_neighbors=15 and min_dist=0.1 were applied for compliance with UMAP assumptions. A composite UMAP projection that utilized data points from all desired samples was generated using Matplotlib. Cells from specific biological samples or timepoints were visualized by overlaying onto the generated UMAP projection which combined all samples and timepoints (https://github.com/lmcinnes/umap).

### Spanning-tree Progression Analysis of Density-normalized Events (SPADE)

SPADE trees generated as previously reported (27, 45, 46). Briefly, SPADE was performed through MATLAB and the source code is available at http://pengqiu.gatech.edu/software/SPADE/. MATLAB-based SPADE automatically generates the tree by performing density-dependent down-sampling, agglomerative clustering, linking clusters with a minimum spanning-tree algorithm and upsampling based on user input. The SPADE tree was generated by exporting uncompensated pregated live, single cells or select pre-gated cellular subsets (i.e. CD3^+^ T cells). The following SPADE parameters were used: Apply compensation matrix in FCS header, Arcsinh transformation with cofactor 150, neighborhood size 5, local density approximation factor 1.5, max allowable cells in pooled downsampled data 50000, target density 20000 cells remaining, and number of desired clusters 50-100, depending on cell population size.

### SPADE node heatmap

SPADE dendrogram heatmaps were constructed with calculated z-scores of fluorescence intensities for each measured surface marker across all nodes of a SPADE dendrogram. Each row of the heatmap corresponds to a surface marker and each column represents a single SPADE node. Marker expression levels range from dark blue to dark red, indicating low to high expression, respectively.

### Quadriceps tissue histology and immunostaining

Tissue processing and histology done as previously reported (24). Briefly, muscle was dissected, weighed, and snap frozen in liquid nitrogen cooled isopentane. 10 μm cryosections (CryoStar NX70 Cryostat) were taken. Samples were blocked and permeabilized before staining with anti-CD206 (Abcam, ab64693, 1:200) for 1hour incubation at room temperature. Secondary antibodies conjugated to Alexa Fluor 555 (Abcam, ab150078, 1:250) and iFluor 488-conjugated phalloidin (Abcam, ab176753, 1:1000) were incubated for 30 mins at room temperature. Slides were mounted with Fluoroshield Mounting Medium with DAPI (Abcam, ab104139) and stored at 4°C. Immunofluorescence images were taken on Zeiss 710 Laser Scanning Confocal microscope at 20x as previously reported (24).

### Statistical Analyses

All statistical analyses were done in GraphPad Prism 8. Data displayed with outlined bars representing the mean, error bars are ± Standard Error of the Mean (S.E.M.). For multiple comparisons, 2-way ANOVA with *Sidak* test was used. To correct for instances of unequal variance and non-normality of cell frequency data, when necessary, statistical analysis was performed on log-10 or square root transformed data, *p*<0.05 considered significant.

## Author Contributions

LAH, SEA, YCJ, NJW, and EAB designed the research studies, analyzed the data, wrote the manuscript, and reviewed the manuscript. LAH, SEA, and TCT performed the experiments, analyzed the data, and reviewed the manuscript. WYY, HSL, and PQ contributed to methodology and reviewed the manuscript.

## Conflict of Interest

The authors have no conflict of interest to declare.

## Funding

This research was supported by the Department of Defense under Award Number W81XWH-20-1-0336 (YCJ). LAH was supported by the National Science Foundation Graduate Research Fellowship (Grant No. DGE-1650044). SEA and TCT were trainees on the NIH/NIGMS-sponsored Cell and Tissue Engineering (CTEng) Biotechnology Training Program (T32GM008433) while this work was conducted.

## Acknowledgements

We thank the core facilities at the Parker H. Petit Institute of Bioengineering and Bioscience at Georgia Institute of Technology for the use of shared equipment, services, and expertise.

## Notes

### Competing Interest Statement

The authors have declared no competing interest.

## References

1. Tidball JG. Regulation of muscle growth and regeneration by the immune system. Nat Rev Immunol. 2017;17(3):165–78. Epub 2017/02/07. doi: 10.1038/nri.2016.150. PubMed PMID: 28163303; PMCID: PMC5452982.

2. Matsuda R, Nishikawa A, Tanaka H. Visualization of Dystrophic Muscle Staining with Evans Blue: Evidence Muscle 1 Fibers in Mdx Mouse of Apoptosis in by Vital. J Biochem. 1995;118:959–64. PubMed PMID: Matsuda1995.

3. Fielding RA, Manfredi TJ, Ding W, Fiatarone MA, Evans WJ, Cannon JG. Acute phase response in exercise III Neutrophil and IL-1p accumu lation in skeletal muscle. Journal of Physiology. 1993(12). PubMed PMID: Fielding1993.

4. Wosczyna MN, Rando TA. A Muscle Stem Cell Support Group: Coordinated Cellular Responses in Muscle Regeneration. Dev Cell. 2018;46(2):135–43. Epub 2018/07/18. doi: 10.1016/j.devcel.2018.06.018. PubMed PMID: 30016618; PMCID: PMC6075730.

5. Joe AW, Yi L, Natarajan A, Le Grand F, So L, Wang J, Rudnicki MA, Rossi FM. Muscle injury activates resident fibro/adipogenic progenitors that facilitate myogenesis. Nat Cell Biol. 2010;12(2):153–63. Epub 2010/01/19. doi: 10.1038/ncb2015. PubMed PMID: 20081841; PMCID: PMC4580288.

6. Lemos DR, Babaeijandaghi F, Low M, Chang CK, Lee ST, Fiore D, Zhang RH, Natarajan A, Nedospasov SA, Rossi FM. Nilotinib reduces muscle fibrosis in chronic muscle injury by promoting TNF-mediated apoptosis of fibro/adipogenic progenitors. Nat Med. 2015;21(7):786–94. Epub 2015/06/09. doi: 10.1038/nm.3869. PubMed PMID: 26053624.

7. Olingy CE, Emeterio CL, Ogle ME, Krieger JR, Bruce AC, Pfau DD, Jordan BT, Peirce SM, Botchwey EA. Non-classical monocytes are biased progenitors of wound healing macrophages during soft tissue injury. Scientific Reports. 7(1). doi: 10.1038/s41598-017-00477-1 PMID - 28348370.

8. Arnold L, Henry A, Poron F, Baba-Amer Y, van Rooijen N, Plonquet A, Gherardi RK, Chazaud B. Inflammatory monocytes recruited after skeletal muscle injury switch into antiinflammatory macrophages to support myogenesis. J Exp Med. 2007;204(5):1057–69. Epub 2007/05/09. doi: 10.1084/jem.20070075. PubMed PMID: 17485518; PMCID: PMC2118577.

9. St Pierre BA, Tidball JG. Differential response of macrophage subpopulations to soleus muscle reloading after rat hindlimb suspension. J Appl Physiol (1985). 1994;77(1):290–7. Epub 1994/07/01. doi: 10.1152/jappl.1994.77.1.290. PubMed PMID: 7961247.

10. Deng B, Wehling-Henricks M, Villalta SA, Wang Y, Tidball JG. IL-10 triggers changes in macrophage phenotype that promote muscle growth and regeneration. J Immunol. 2012;189(7):3669–80. Epub 2012/08/31. doi: 10.4049/jimmunol.1103180. PubMed PMID: 22933625; PMCID: PMC3448810.

11. Chazaud B. Inflammation and Skeletal Muscle Regeneration: Leave It to the Macrophages! Trends Immunol. 2020;41(6):481–92. Epub 2020/05/05. doi: 10.1016/j.it.2020.04.006. PubMed PMID: 32362490.

12. Wang H, Melton DW, Porter L, Sarwar ZU, McManus LM, Shireman PK. Altered macrophage phenotype transition impairs skeletal muscle regeneration. Am J Pathol. 2014;184(4):1167–84. Epub 2014/02/15. doi: 10.1016/j.ajpath.2013.12.020. PubMed PMID: 24525152; PMCID: PMC3969996.

13. Summan M, Warren GL, Mercer RR, Chapman R, Hulderman T, Van Rooijen N, Simeonova PP. Macrophages and skeletal muscle regeneration: a clodronate-containing liposome depletion study. Am J Physiol Regul Integr Comp Physiol. 2006;290(6):R1488–95. Epub 2006/01/21. doi: 10.1152/ajpregu.00465.2005. PubMed PMID: 16424086.

14. Zhang J, Xiao Z, Qu C, Cui W, Wang X, Du J. CD8 T cells are involved in skeletal muscle regeneration through facilitating MCP-1 secretion and Gr1(high) macrophage infiltration. J Immunol. 2014;193(10):5149–60. Epub 2014/10/24. doi: 10.4049/jimmunol.1303486. PubMed PMID: 25339660.

15. Burzyn D, Kuswanto W, Kolodin D, Shadrach JL, Cerletti M, Jang Y, Sefik E, Tan TG, Wagers AJ, Benoist C, Mathis D. A special population of regulatory T cells potentiates muscle repair. Cell. 2013;155(6):1282–95. Epub 2013/12/10. doi: 10.1016/j.cell.2013.10.054. PubMed PMID: 24315098; PMCID: PMC3894749.

16. Villalta SA, Rosenthal W, Martinez L, Kaur A, Sparwasser T, Tidball JG, Margeta M, Spencer MJ, Bluestone JA. Regulatory T cells suppress muscle inflammation and injury in muscular dystrophy. Sci Transl Med. 2014;6(258):258ra142. Epub 2014/10/17. doi: 10.1126/scitranslmed.3009925. PubMed PMID: 25320234; PMCID: PMC4889432.

17. Panduro M, Benoist C, Mathis D. Treg cells limit IFN-gamma production to control macrophage accrual and phenotype during skeletal muscle regeneration. Proc Natl Acad Sci U S A. 2018;115(11):E2585–E93. Epub 2018/02/25. doi: 10.1073/pnas.1800618115. PubMed PMID: 29476012; PMCID: PMC5856564.

18. Grogan BF, Hsu JR, Skeletal Trauma Research C. Volumetric muscle loss. J Am Acad Orthop Surg. 2011;19 Suppl 1:S35–7. Epub 2011/03/17. doi: 10.5435/00124635-201102001-00007. PubMed PMID: 21304045.

19. Corona BT, Rivera JC, Owens JG, Wenke JC, Rathbone CR. Volumetric muscle loss leads to permanent disability following extremity trauma. J Rehabil Res Dev. 2015;52(7):785–92. Epub 2016/01/09. doi: 10.1682/JRRD.2014.07.0165. PubMed PMID: 26745661.

20. Pollot BEC, B.T. In: Kyba M, editor. Skeletal Muscle Regeneration in the Mouse Methods and Protocols: Springer; 2017. p. 1–14.

21. Novak ML, Weinheimer-Haus EM, Koh TJ. Macrophage activation and skeletal muscle healing following traumatic injury. J Pathol. 2014;232(3):344–55. Epub 2013/11/21. doi: 10.1002/path.4301. PubMed PMID: 24255005; PMCID: PMC4019602.

22. Kuswanto W, Burzyn D, Panduro M, Wang KK, Jang YC, Wagers AJ, Benoist C, Mathis D. Poor Repair of Skeletal Muscle in Aging Mice Reflects a Defect in Local, Interleukin-33-Dependent Accumulation of Regulatory T Cells. Immunity. 2016;44(2):355–67. Epub 2016/02/14. doi: 10.1016/j.immuni.2016.01.009. PubMed PMID: 26872699; PMCID: PMC4764071.

23. Contreras O, Cruz-Soca M, Theret M, Soliman H, Tung LW, Groppa E, Rossi FM, Brandan E. Cross-talk between TGF-beta and PDGFRalpha signaling pathways regulates the fate of stromal fibro-adipogenic progenitors. J Cell Sci. 2019;132(19). Epub 2019/08/23. doi: 10.1242/jcs.232157. PubMed PMID: 31434718.

24. Anderson SE, Han WM, Srinivasa V, Mohiuddin M, Ruehle MA, Moon JY, Shin E, San Emeterio CL, Ogle ME, Botchwey EA, Willett NJ, Jang YC. Determination of a Critical Size Threshold for Volumetric Muscle Loss in the Mouse Quadriceps. Tissue Eng Part C Methods. 2019;25(2):59–70. Epub 2019/01/17. doi: 10.1089/ten.TEC.2018.0324. PubMed PMID: 30648479; PMCID: PMC6389771.

25. Trapnell C, Cacchiarelli D, Grimsby J, Pokharel P, Li S, Morse M, Lennon NJ, Livak KJ, Mikkelsen TS, Rinn JL. The dynamics and regulators of cell fate decisions are revealed by pseudotemporal ordering of single cells. Nat Biotechnol. 2014;32(4):381–6. Epub 2014/03/25. doi: 10.1038/nbt.2859. PubMed PMID: 24658644; PMCID: PMC4122333.

26. Ramskold D, Luo S, Wang YC, Li R, Deng Q, Faridani OR, Daniels GA, Khrebtukova I, Loring JF, Laurent LC, Schroth GP, Sandberg R. Author Correction: Full-length mRNA-Seq from single-cell levels of RNA and individual circulating tumor cells. Nat Biotechnol. 2020;38(3):374. Epub 2020/02/06. doi: 10.1038/s41587-020-0427-1. PubMed PMID: 32015550.

27. Turner TC, Sok MCP, Hymel LA, Pittman FS, York WY, Mac QD, Vyshnya S, Lim HS, Kwong GA, Qiu P, Botchwey EA. Harnessing lipid signaling pathways to target specialized pro-angiogenic neutrophil subsets for regenerative immunotherapy. Sci Adv. 2020;6(44). Epub 2020/11/01. doi: 10.1126/sciadv.aba7702. PubMed PMID: 33127670; PMCID: PMC7608810.

28. Becht E, McInnes L, Healy J, Dutertre CA, Kwok IWH, Ng LG, Ginhoux F, Newell EW. Dimensionality reduction for visualizing single-cell data using UMAP. Nat Biotechnol. 2018. Epub 2018/12/12. doi: 10.1038/nbt.4314. PubMed PMID: 30531897.

29. Anchang B, Hart TD, Bendall SC, Qiu P, Bjornson Z, Linderman M, Nolan GP, Plevritis SK. Visualization and cellular hierarchy inference of single-cell data using SPADE. Nat Protoc. 2016;11(7):1264–79. Epub 2016/06/17. doi: 10.1038/nprot.2016.066. PubMed PMID: 27310265.

30. Qiu P, Simonds EF, Bendall SC, Gibbs KD, Jr., Bruggner RV, Linderman MD, Sachs K, Nolan GP, Plevritis SK. Extracting a cellular hierarchy from high-dimensional cytometry data with SPADE. Nat Biotechnol. 2011;29(10):886–91. Epub 2011/10/04. doi: 10.1038/nbt.1991. PubMed PMID: 21964415; PMCID: PMC3196363.

31. Braga TT, Agudelo JS, Camara NO. Macrophages During the Fibrotic Process: M2 as Friend and Foe. Front Immunol. 2015;6:602. Epub 2015/12/05. doi: 10.3389/fimmu.2015.00602. PubMed PMID: 26635814; PMCID: PMC4658431.

32. Aguilar CA, Greising SM, Watts A, Goldman SM, Peragallo C, Zook C, Larouche J, Corona BT. Correction: Multiscale analysis of a regenerative therapy for treatment of volumetric muscle loss injury. Cell Death Discov. 2018;4:16. Epub 2018/08/01. doi: 10.1038/s41420-018-0080-3. PubMed PMID: 30062061; PMCID: PMC6056478.

33. Palit S, Heuser C, de Almeida GP, Theis FJ, Zielinski CE. Meeting the Challenges of HighDimensional Single-Cell Data Analysis in Immunology. Front Immunol. 2019;10:1515. Epub 2019/07/30. doi: 10.3389/fimmu.2019.01515. PubMed PMID: 31354705; PMCID: PMC6634245.

34. San Emeterio CL, Olingy CE, Chu Y, Botchwey EA. Selective recruitment of non-classical monocytes promotes skeletal muscle repair. Biomaterials. 2017;117:32–43. Epub 2016/12/09. doi: 10.1016/j.biomaterials.2016.11.021. PubMed PMID: 27930948; PMCID: PMC5218730.

35. Chen Y, Pu Q, Ma Y, Zhang H, Ye T, Zhao C, Huang X, Ren Y, Qiao L, Liu HM, Esmon CT, Ding BS, Cao Z. Aging Reprograms the Hematopoietic-Vascular Niche to Impede Regeneration and Promote Fibrosis. Cell Metab. 2021;33(2):395–410 e4. Epub 2020/12/29. doi: 10.1016/j.cmet.2020.11.019. PubMed PMID: 33357457.

36. Borthwick LA, Wynn TA, Fisher AJ. Cytokine mediated tissue fibrosis. Biochim Biophys Acta. 2013;1832(7):1049–60. Epub 2012/10/11. doi: 10.1016/j.bbadis.2012.09.014. PubMed PMID: 23046809; PMCID: PMC3787896.

37. Villalta SA, Nguyen HX, Deng B, Gotoh T, Tidball JG. Shifts in macrophage phenotypes and macrophage competition for arginine metabolism affect the severity of muscle pathology in muscular dystrophy. Hum Mol Genet. 2009;18(3):482–96. Epub 2008/11/11. doi: 10.1093/hmg/ddn376. PubMed PMID: 18996917; PMCID: PMC2638796.

38. Lech M, Anders HJ. Macrophages and fibrosis: How resident and infiltrating mononuclear phagocytes orchestrate all phases of tissue injury and repair. Biochim Biophys Acta. 2013;1832(7):989–97. Epub 2012/12/19. doi: 10.1016/j.bbadis.2012.12.001. PubMed PMID: 23246690.

39. Guerriero JL. Macrophages: Their Untold Story in T Cell Activation and Function. Int Rev Cell Mol Biol. 2019;342:73–93. Epub 2019/01/13. doi: 10.1016/bs.ircmb.2018.07.001. PubMed PMID: 30635094.

40. Bohmer RM, Bandala-Sanchez E, Harrison LC. Forward light scatter is a simple measure of T-cell activation and proliferation but is not universally suited for doublet discrimination. Cytometry A. 2011;79(8):646–52. Epub 2011/06/23. doi: 10.1002/cyto.a.21096. PubMed PMID: 21695774.

41. Overgaard NH, Jung JW, Steptoe RJ, Wells JW. CD4+/CD8+ double-positive T cells: more than just a developmental stage? J Leukoc Biol. 2015;97(1):31–8. Epub 2014/11/02. doi: 10.1189/jlb.1RU0814-382. PubMed PMID: 25360000.

42. Simonetta F, Chiali A, Cordier C, Urrutia A, Girault I, Bloquet S, Tanchot C, Bourgeois C. Increased CD127 expression on activated FOXP3+CD4+ regulatory T cells. Eur J Immunol. 2010;40(9):2528–38. Epub 2010/08/07. doi: 10.1002/eji.201040531. PubMed PMID: 20690182.

43. Haugen F, Norheim F, Lian H, Wensaas AJ, Dueland S, Berg O, Funderud A, Skalhegg BS, Raastad T, Drevon CA. IL-7 is expressed and secreted by human skeletal muscle cells. Am J Physiol Cell Physiol. 2010;298(4):C807–16. Epub 2010/01/22. doi: 10.1152/ajpcell.00094.2009. PubMed PMID: 20089933.

44. Dejaco C, Duftner C, Grubeck-Loebenstein B, Schirmer M. Imbalance of regulatory T cells in human autoimmune diseases. Immunology. 2006;117(3):289–300. Epub 2006/02/16. doi: 10.1111/j.1365-2567.2005.02317.x. PubMed PMID: 16476048; PMCID: PMC1782226.

45. Hymel LA, Ogle ME, Anderson SE, San Emeterio CL, Turner TC, York WY, Liu AY, Olingy CE, Sridhar S, Lim HS, Sulchek T, Qiu P, Jang YC, Willett NJ, Botchwey EA. Modulating local S1P receptor signaling as a regenerative immunotherapy after volumetric muscle loss injury. J Biomed Mater Res A. 2021;109(5):695–712. Epub 2020/07/02. doi: 10.1002/jbm.a.37053. PubMed PMID: 32608188; PMCID: PMC7772280.

46. San Emeterio CL, Hymel LA, Turner TC, Ogle ME, Pendleton EG, York WY, Olingy CE, Liu AY, Lim HS, Sulchek TA, Warren GL, Mortensen LJ, Qiu P, Jang YC, Willett NJ, Botchwey EA. Nanofiber-Based Delivery of Bioactive Lipids Promotes Pro-regenerative Inflammation and Enhances Muscle Fiber Growth After Volumetric Muscle Loss. Front Bioeng Biotechnol. 2021;9:650289. Epub 2021/04/06. doi: 10.3389/fbioe.2021.650289. PubMed PMID: 33816455; PMCID: PMC8017294.

