## Supplementary material for "Identifying dysregulated immune cell subsets following critical volumetric muscle loss with pseudo-time trajectories": Figure S1, Figure S2, Figure S3, Figure S4, Figure S5

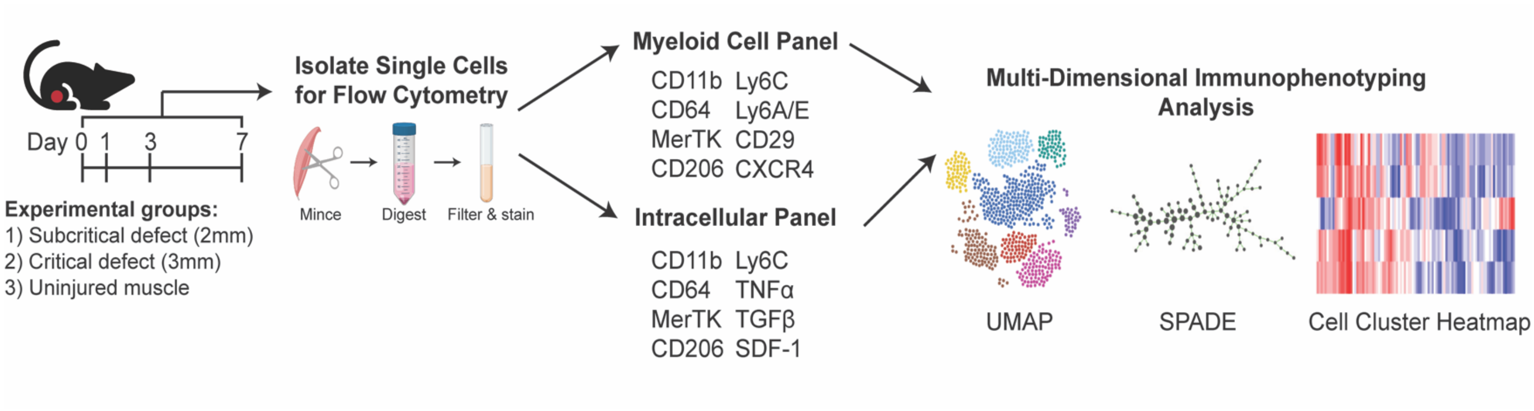


**Supplementary Figure 1. Graphical workflow of multiparameter pseudo-time analysis applied to single cell flow cytometry data.** Animals received either a full-thickness subcritical (2 mm diameter biopsy) or critical (3 mm diameter biopsy) unilateral VML injury to the left quadriceps. Uninjured quadriceps muscle was used as a control. At 1, 3, or 7 days post injury, tissue was excised, digested, and single cells were isolated and stained for flow cytometric analysis. Dimensionality reduction and pseudo-time clustering algorithms, namely UMAP and SPADE, were employed to characterize the temporal immune response of myeloid and lymphoid cells following VML.

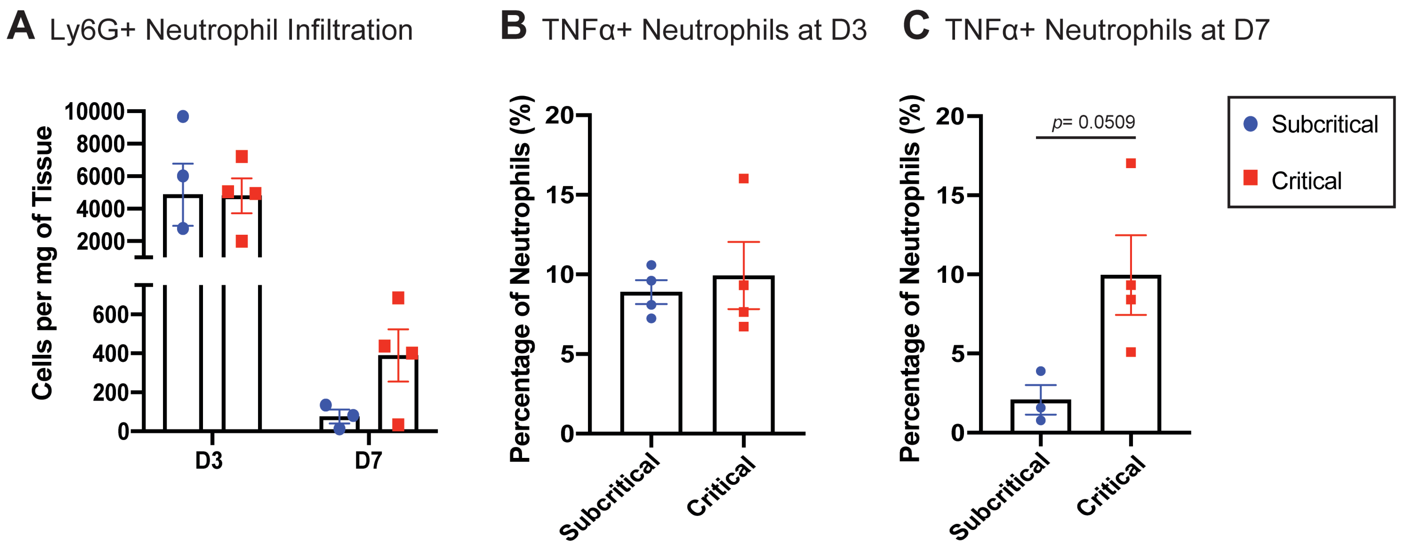


**Supplementary Figure 2. TNF-α+ neutrophils persist within critically sized VML injuries at day 7.** Single cell flow cytometric analysis performed at days 3 and 7 post subcritical or critical VML injury to murine quadriceps. **(A)** CD11b+Ly6G+ neutrophil concentration within injured quadriceps at each timepoint post injury. **(B)** The percentage of total neutrophils that express TNF-α from subcritical and critical VML injuries at day 3. **(C)** The percentage of total neutrophils that express TNF-α from subcritical and critical VML injuries at day 7. Data presented as mean ± S.E.M. Statistical analysis conducted with unpaired t-test (C). n= 3-4 animals per experimental group. D3: day 3, D7: day 7.

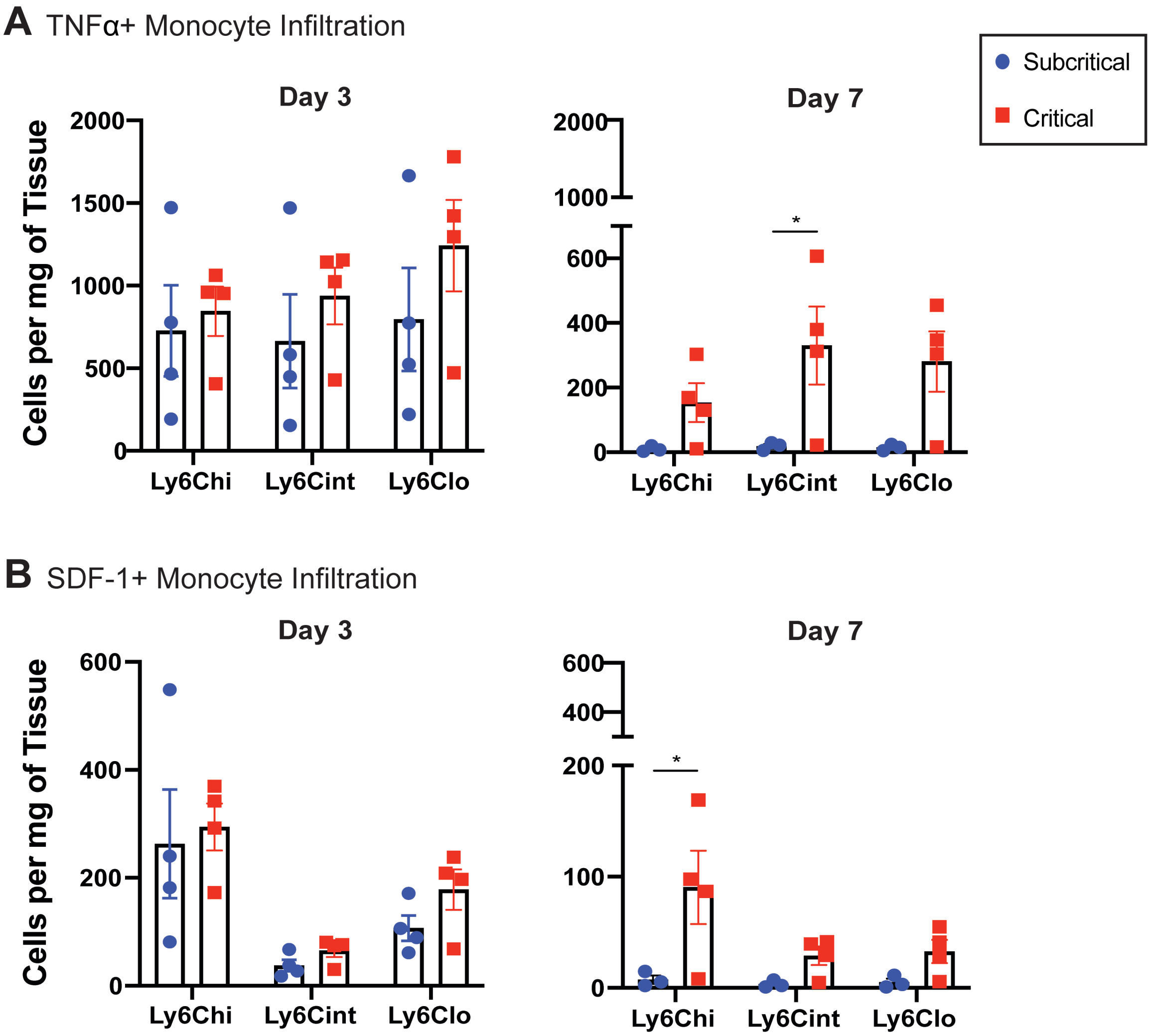


**Supplementary Figure 3. Monocyte subsets infiltrating critically sized defects may propagate chronic inflammation via TNF-α and SDF-1 cytokine signaling.** Single cell flow cytometric analysis performed at days 3 and 7 post subcritical or critical VML injury to murine quadriceps. $CD11b^{+}CD64^{+}MerTK^{-}SSC^{lo}$ monocytes characterized into $Ly6C^{lo}$, $Ly6C^{int}$, or $Ly6C^{hi}$ subsets based on expression of Ly6C. **(A)** Concentration of each monocyte subset expressing TNF-α at day 3 and 7 post subcritical or critical VML injury. **(B)** Concentration of each monocyte subset expressing SDF-1 at day 3 and 7 post subcritical or critical VML injury. Data presented as mean ± S.E.M. Statistical analyses include two-way ANOVA with *Sidak* test for multiple comparisons between injury sizes. *$p$<0.05, n= 3-4 animals per experimental group.

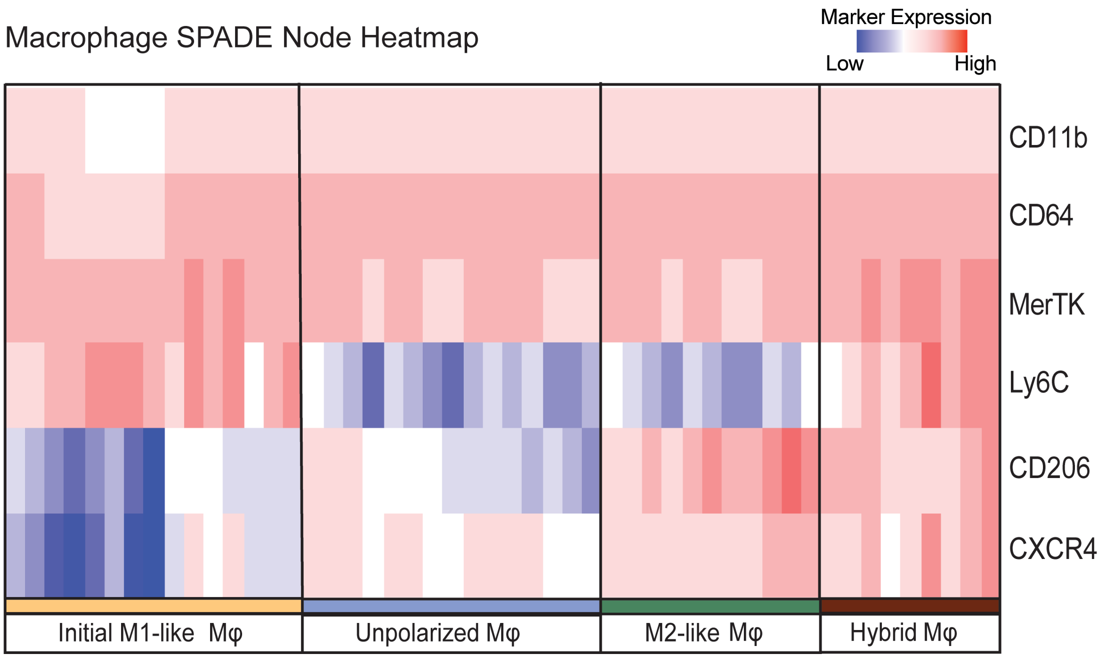


**Supplementary Figure 4. Characterization of macrophage subset phenotypes visualized by SPADE node marker expression.** Heatmap rendering of marker expression levels within each macrophage SPADE node ranging from dark blue to dark red, representing low to high marker expression, respectively. Each column of the heatmap represents one node and each row is one surface marker as labeled. SPADE dendrogram was comprised of $CD11b^{+}CD64^{+}MerTK^{+}$ macrophages pooled from all injury groups (uninjured, subcritical, and critical VML) and timepoints (days 1, 3, and 7). M1-like macrophage nodes were identified by high expression of Ly6C and low expression of CD206. Unpolarized macrophages were characterized by the dual low expression of CD206 and Ly6C. M2-like macrophages present with high CD206 expression and low Ly6C, and hybrid macrophages express both CD206 and Ly6C together.

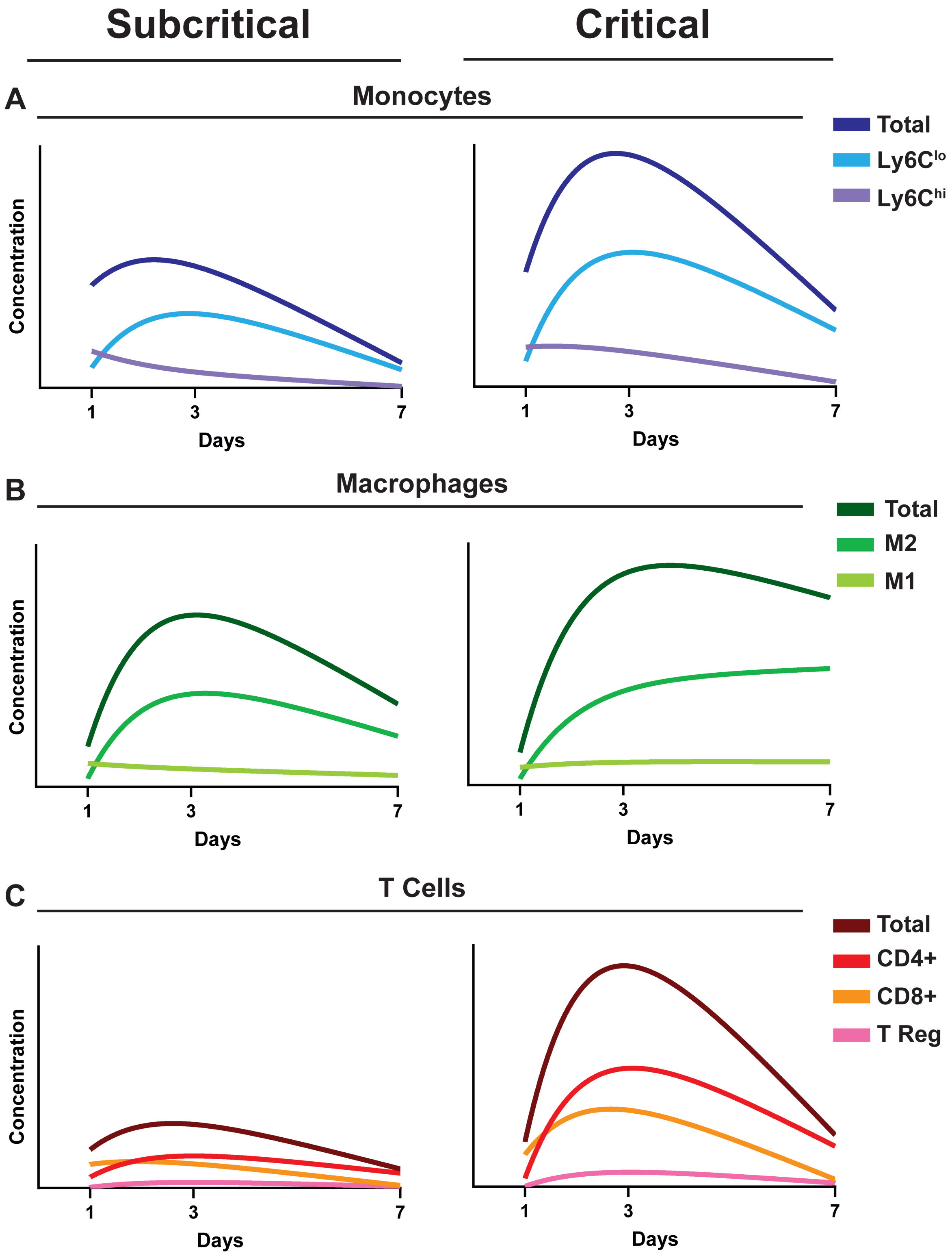


**Supplementary Figure 5. Temporal evaluation of immune cell response to subcritical and critical VML injury.** Summary of the immune cell response in the first week following subcritical and critical VML injuries. Cell population dynamics from each injury size represented as lines fit with a continuous hinge function (GraphPad Prism 8) through the quantified concentration data for monocytes **(A)**, macrophages **(B)**, and T cells **(C)**. Axes values were kept consistent between cell populations.
